# Structural characterisation of KKT4, an unconventional microtubule-binding kinetochore protein

**DOI:** 10.1101/2020.10.14.337170

**Authors:** Patryk Ludzia, Edward D. Lowe, Gabriele Marcianò, Shabaz Mohammed, Christina Redfield, Bungo Akiyoshi

## Abstract

The kinetochore is the macromolecular protein machinery that drives chromosome segregation by interacting with spindle microtubules. Unlike most other eukaryotes that have canonical kinetochore proteins, a group of evolutionarily divergent eukaryotes called kinetoplastids (such as *Trypanosoma brucei*) have a unique set of kinetochore proteins. To date, KKT4 is the only kinetoplastid kinetochore protein that is known to bind microtubules. Here we use X-ray crystallography, NMR spectroscopy, and crosslinking mass spectrometry to characterise the structure and dynamics of KKT4. We show that its microtubule-binding domain consists of a coiled-coil structure followed by a positively charged disordered tail. The crystal structure of the C-terminal BRCT domain of KKT4 reveals that it is likely a phosphorylation-dependent protein-protein interaction domain. The BRCT domain interacts with the N-terminal region of the KKT4 microtubule-binding domain and with a phosphopeptide derived from KKT8. Finally, we show that KKT4 binds DNA with high affinity. Taken together, these results provide the first structural insights into the unconventional kinetoplastid kinetochore protein KKT4.

## Introduction

Every time a cell divides, it must duplicate and segregate its genetic material accurately into two daughter cells. A key structure involved in chromosome segregation in eukaryotes is the kinetochore, a macromolecular protein complex that assembles onto centromeric DNA and interacts with spindle microtubules during mitosis and meiosis (McIntosh, 2016). Microtubules are dynamic polymers that change in length by addition or removal of tubulin subunits at the tips (Desai and Mitchison, 1997). Accurate chromosome segregation requires that kinetochores form robust attachments to the dynamic microtubule tips. In addition, kinetochores need to destabilise erroneous attachments to ensure that sister kinetochores bind microtubules emanating from opposite poles (Nicklas, 1997, Biggins, 2013, Cheeseman, 2014, Musacchio and Desai, 2017). Revealing the molecular basis of kinetochore-microtubule attachments and their regulation is key to understanding the mechanism of chromosome segregation.

Studies on commonly-studied eukaryotes have identified a number of microtubule-binding kinetochore proteins, including the Ndc80, Dam1, Ska, and SKAP-Astrin complexes (Cheeseman et al., 2001, Cheeseman et al., 2006, Hanisch et al., 2006, Schmidt et al., 2012, Abad et al., 2014, Friese et al., 2016, Kern et al., 2017). Some of these components are widely conserved among eukaryotes (Meraldi et al., 2006, van Hooff et al., 2017). However, none of these or other canonical structural kinetochore components has been identified in kinetoplastids, an evolutionarily-divergent group of unicellular flagellated eukaryotes such as parasitic trypanosomatids (e.g. *Trypanosoma brucei*, *Trypanosoma cruzi*, and *Leishmania* species) and free-living bodonids (e.g. *Bodo saltans*) (Berriman et al., 2005, Cavalier-Smith, 2010). Instead, a number of unique kinetochore proteins have been identified in *T. brucei,* namely 24 kinetoplastid kinetochore proteins (KKT1–20, KKT22–25) and 12 KKT-interacting proteins (KKIP1–12) (Akiyoshi and Gull, 2014, Nerusheva and Akiyoshi, 2016, D’Archivio and Wickstead, 2017, Brusini et al., 2019, Nerusheva et al., 2019). These proteins do not appear orthologous to canonical kinetochore proteins in other eukaryotes, suggesting that kinetoplastids use a distinct set of proteins to build up unique kinetochores. We previously identified KKT4 as the first microtubule-binding kinetochore protein in *T. brucei* (throughout this manuscript we refer to KKT4 from *Trypanosoma brucei* unless stated otherwise) (Llauró et al., 2018). KKT4 directly binds to microtubules and maintains load-bearing attachments to both growing and shortening microtubule tips *in vitro*. Microtubule-binding activities were also found in KKT4 from other kinetoplastids (*T. cruzi*, *T. congolense, Leishmania mexicana*, and *Phytomonas*), suggesting that KKT4 plays an important role in kinetochore-microtubule attachments in kinetoplastids (Llauró et al., 2018). Using microtubule co-sedimentation assays, we defined KKT4^115–343^ as the microtubule-binding domain in *T. brucei*. Importantly, KKT4^115–343^ lacks significant sequence similarity to the microtubule-binding domain of any known kinetochore protein (such as the Ndc80, Dam1, and Ska complexes). To date, there is no structural information available for KKT4. It therefore remains unknown how KKT4 forms attachments to microtubules and whether its microtubule-binding activity is regulated. Interestingly, KKT4 has a putative BRCA1 C-terminal (BRCT) domain, which is not present in any known kinetochore protein in other eukaryotes (Figure 1A) (Akiyoshi and Gull, 2014). The function of this putative BRCT domain also remains unknown.

**Figure 1.**
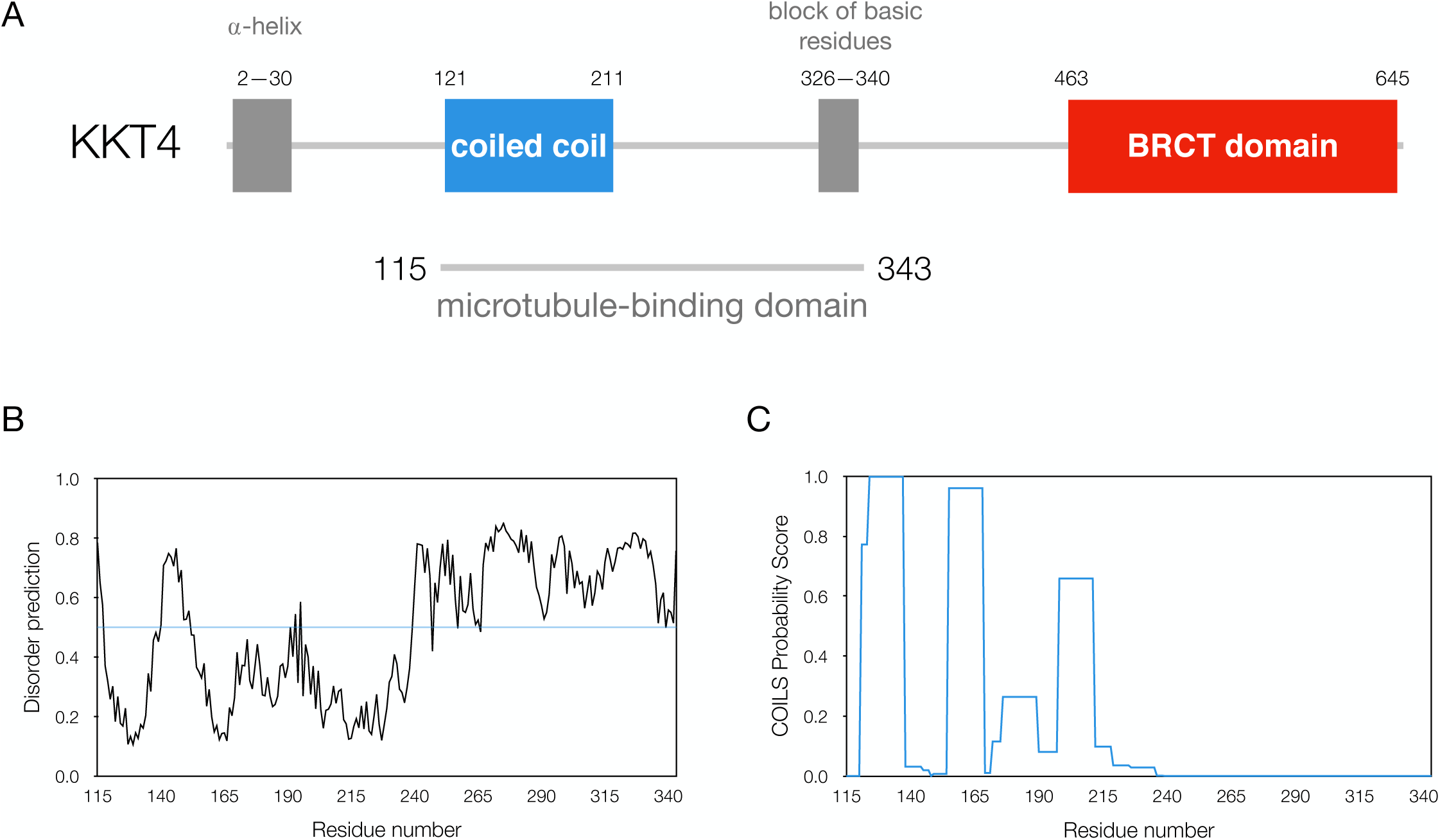
Domain organisation and structure predictions for *T. brucei* KKT4. (A) KKT4 has the following regions conserved among kinetoplastid species: an N-terminal predicted α-helix, predicted coiled-coil region, block of basic residues, and a putative BRCT domain at the C-terminus. (B) Disorder prediction for KKT4^115–343^ using DisEMBL (Linding et al., 2003). The N-terminus of the microtubule-binding domain is predicted to be mainly ordered (118–239), while the C-terminus is predicted to be mostly disordered. The blue horizontal line at 0.5 indicates the boundary between predicted order and disorder. (C)Coiled coil prediction of KKT4^115–343^ by COILS (window = 14) (Lupas et al., 1991). Coiled coils are predicted from 121 to 211 with breaks between 138–154 and 169–189.

Here, we have used X-ray crystallography and NMR spectroscopy to obtain structural information on the KKT4 protein. Its microtubule-binding domain consists of a coiled-coil structure followed by a positively charged disordered tail. A crystal structure of the C-terminal BRCT domain reveals a putative phosphopeptide-binding pocket. Moreover, crosslinking mass spectrometry (XL-MS) and NMR have revealed interactions between the microtubule-binding domain and the BRCT domain. Overall these analyses show that the KKT4 structure is distinct from any known microtubule-binding kinetochore protein.

## Results

### KKT4 forms oligomers

Our previous single-molecule experiments suggested that KKT4 has a tendency to oligomerise even at a low nanomolar concentration (Llauró et al., 2018). To determine the oligomerisation state of KKT4 in solution, we used size exclusion chromatography coupled with multi-angle light scattering (SEC-MALS) (Wen et al., 1996) (Table 1 and Supplemental Figure 1). Analysis of the full-length protein revealed that KKT4 is tetrameric under the conditions used (Supplemental Figure 1A). To identify the region(s) responsible for the oligomerisation, smaller fragments of KKT4 were analysed. The following regions of KKT4 are well conserved among kinetoplastids: N-terminal predicted *α*-helices, predicted coiled coils within the microtubule-binding domain, a block of basic residues, and the C-terminal BRCT domain (Figure 1A) (Llauró et al., 2018). We found that KKT4^463– 645^, that has a putative BRCT domain, behaved as a monomer (Supplemental Figure 1B), while KKT4^101–352^ that contains the microtubule-binding domain behaved as a dimer (Supplemental Figure 1C). In contrast, KKT4^2–114^ showed characteristics of a tetramer at higher concentrations but of a trimer at lower concentrations (Supplemental Figure 1D). Thus, the N-terminal region is responsible for the formation of the KKT4 tetramer. It is possible that KKT4^2–114^ is in a dimer-tetramer equilibrium (rather than trimer-tetramer) because the microtubule-binding domain is a dimer and the full-length protein is likely to be a dimer of dimers (Supplemental Figure 1A). These results suggest that KKT4 has multiple regions that promote oligomerisation.

**Table 1.** SEC-MALS results for KKT4 fragments

| Protein | Concentration<br>[μM] | Monomer<br>molecular weight<br>[kDa] | Observed<br>molecular weight<br>[kDa] | Estimated<br>oligomerisation<br>state |
| --- | --- | --- | --- | --- |
| SNAP-6HIS-3FLAG-KKT4<sup>2-645</sup> | 20 | 92 | 350 | Tetramer |
| KKT4<sup>463-645</sup> | 100 | 20 | 19 | Monomer |
| KKT4<sup>101-352</sup> | 35 | 28 | 55 | Dimer |
| KKT4<sup>2-114</sup> | 1000/800/500/<br>350/170 | 11 | 37.9/34.6/31.5/<br>32.8/28.1 | Tetramer-trimer or<br>tetramer-dimer<br>mixture |

### The C-terminal part of the KKT4 microtubule-binding domain is disordered

To gain structural insights into the microtubule-binding domain of KKT4, we tried to crystallise KKT4^115–343^. Despite extensive attempts, however, no suitable crystals were obtained. It is likely that the predicted disordered region in the C-terminal part of KKT4^115–343^ prevented the formation of diffraction quality crystals (Figure 1B). We therefore employed NMR spectroscopy to probe the structure and dynamics of the KKT4 microtubule-binding domain in solution. The 2D ^1^H-^15^N correlation spectrum of ^15^N-KKT4^115–343^ showed a large variation in peak intensities (Supplemental Figure 2A); this suggests a mixture of structured and disordered regions in KKT4^115–343^. The strongest peaks in the spectrum of KKT4^115–343^ were assigned to residues between 115 and 118 in the N-terminus and those between 231 and 343 in the C-terminal half of the fragment (Ludzia et al., 2020). Weaker peaks have been assigned to residues between 119 and 150, while no peaks were observed for residues 151 to 230 (Ludzia et al., 2020).

Backbone ^1^H, ^13^C and ^15^N chemical shifts are sensitive indicators of secondary structure (Spera and Bax, 1991, Wishart et al., 1991, Beger and Bolton, 1997, Cornilescu et al., 1999). Analysis of ^1^H*α*, ^1^HN, ^13^C*α*, ^13^Cβ, ^13^CO and ^15^N chemical shifts using TALOS-N (Shen and Bax, 2013) predicted no stable secondary structure for residues 231 to 343 (data not shown). The Secondary Structure Propensity (SSP) score, which is more suitable in identifying structural propensities for disordered proteins (Marsh et al., 2006), also found no secondary structure propensity greater than 0.25 (Figure 2A), suggesting that the C-terminus of KKT4^115–343^ from residue 231 onwards is indeed disordered.

**Figure 2.**
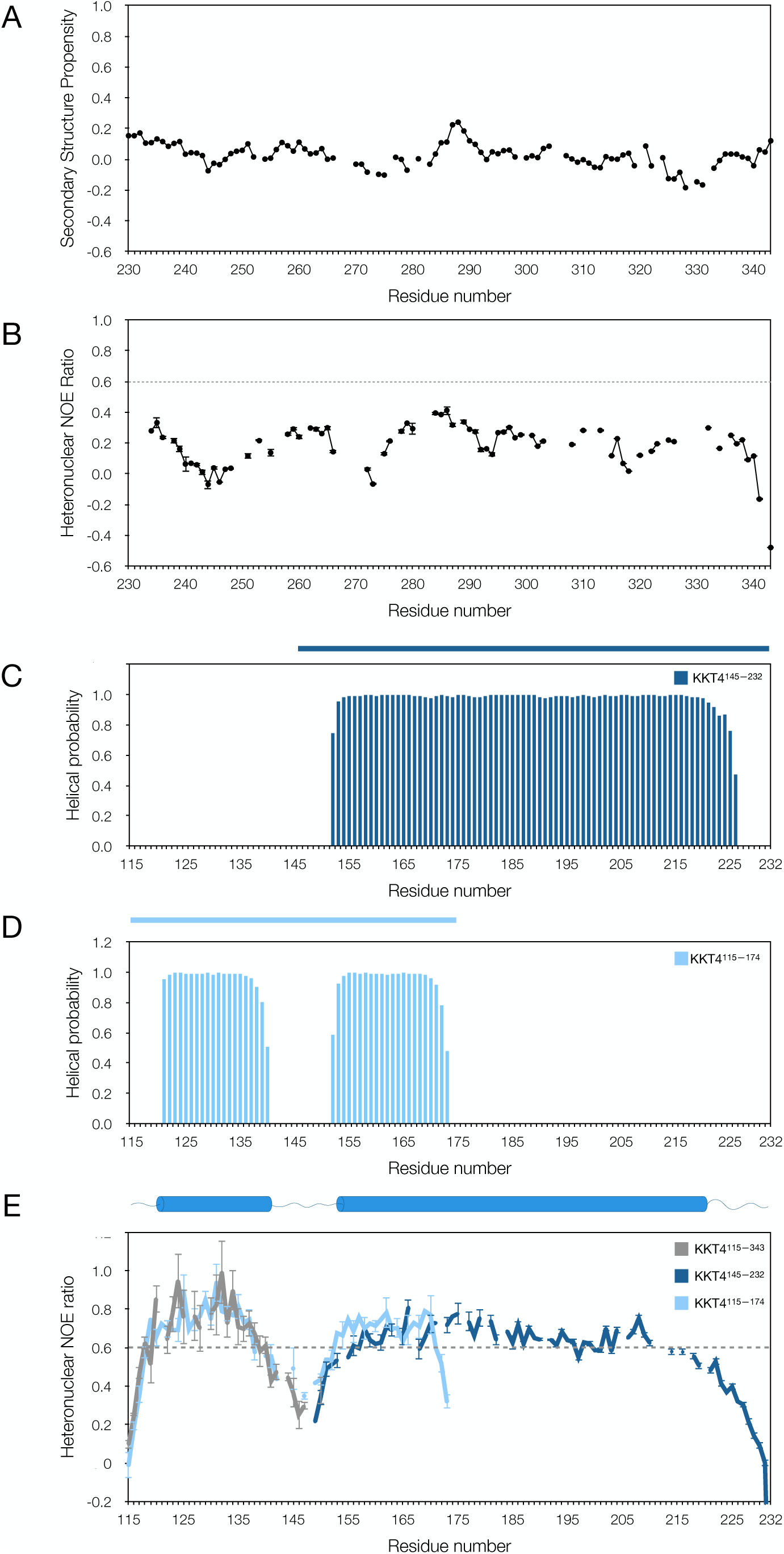
NMR analysis of the KKT4 microtubule-binding domain. (A) Secondary structure propensity (SSP) scores (Marsh et al., 2006) for residues 230–343 of KKT4^115–343^ plotted against the protein sequence show no secondary structure propensity of greater than 0.25. The gaps in the graph correspond to residues that could not be assigned. (B) The {^1^H}-^15^N heteronuclear NOE ratio plotted as a function of the KKT4^115–343^ sequence has values of less than 0.6 for all residues between 230 and 343, showing that this region is disordered. Variation of the NOE is observed along the sequence indicating some variability in the degree of flexibility. The reduced values at ∼245 coincide with a Gly-Gly-Gly sequence at 242–244 likely due to the absence of a sidechain for glycine that allows this residue to sample a wider range of backbone conformations. Lower NOE ratios are also observed in the vicinity of the Gly-Ser-Gly sequence at 271–273. However, not all glycine residues are located in the most flexible regions. G285, which has an NOE ratio of 0.38, is located in a region with the highest NOE ratios, consistent with the SSP prediction. (C)TALOS-N secondary structure prediction for KKT4^145–232^ shows a continuous helix from 152 to 225. The dark blue bar above the graph indicates the length of KKT4 fragment used for data acquisition and TALOS-N analysis. (D) TALOS-N secondary structure prediction for KKT4^115–174^ shows two regions of helical structure separated by an unstructured linker. The first helix encompasses residues 121–139 and the second helix encompasses residues 152–172. The light blue bar above the graph indicates the length of KKT4 fragment used for data collection and TALOS-N analysis. (E) The {^1^H}-^15^N heteronuclear NOE ratio plotted as a function of sequence for residues 115–232 (values for the C-terminus of KKT4^115–343^ are shown in panel B). Ratios measured for ^15^N-KKT4^115–343^ are shown in grey, ^15^N-KKT4^145–232^ in dark blue, and ^15^N-KKT4^115–174^ in light blue. The heteronuclear NOE values in the region of the 1^st^ helix for KKT4^115–343^ and KKT4^115–174^ agree well, suggesting that removal of the disordered C-terminus did not affect the properties of the KKT4^115–174^ N-terminus. The lower ratios at the beginnings and ends of the helices may indicate helix fraying; this is consistent with the SSP predictions which gave helical propensities of 0.5–0.8 for N- and C-terminal residues of the helices but values of > 0.8 in between (Supplemental Figure 4) (Marsh et al., 2006). The C-terminus of KKT4^115–174^ has significantly lower heteronuclear NOE ratios compared to the same residues in KKT4^145–232^ because G174 is the artificially designed C-terminus of KKT4^115–174^. The value for the C-terminal residue Y232 (−1.16) in KKT4^145–232^ is not shown for clarity of the figure. The top panel shows a summary cartoon of the overall secondary-structure of KKT4^115–232^ predicted by TALOS-N.

To probe the dynamics of the C-terminus of KKT4^115–343^, the {^1^H}–^15^N heteronuclear NOE, which is sensitive to backbone motions on a timescale (ps) faster than the overall tumbling of the molecule (ns), was measured (Figure 2B). A {^1^H}–^15^N NOE ratio of greater than ∼0.6 is characteristic of a structured backbone, while values below ∼0.6 indicate higher flexibility (Kay et al., 1989). For KKT4^115–343^, a {^1^H}–^15^N NOE ratio of less than 0.6 was found for all residues from 231 to 343 (Figure 2B). Taken together, the chemical shift analysis and heteronuclear NOE confirmed disorder in the C-terminal part of KKT4^115–343^.

### The N-terminal part of the KKT4 microtubule-binding domain is structured

The N-terminal region of KKT4^115–343^ is predicted to adopt a coiled-coil structure (Figure 1C). Information for residues preceding 231 could not be obtained from the NMR spectra of KKT4^115–343^ due to weak or missing peaks for these residues. In order to gain insights into the structure and dynamics of this region, NMR data were collected for a shorter fragment (KKT4^115–232^) lacking the flexible C-terminal region. Comparison of the spectra of KKT4^115–232^ and KKT4^115–343^ showed that they share the same group of weak peaks coming from the N-terminal residues (Supplemental Figure 2B), indicating that the removal of the C-terminal disordered tail did not perturb the structure between residues 115 and 150. However, the peaks from residues 151 to 221 were still missing from the spectrum of KKT4^115–232^. The absence of these peaks could be due to the elongated structure of a coiled coil, which would tumble in a non-uniform way and result in broad ^1^H^N^-^15^N signals in helices (Mackay et al., 1996, Schnell et al., 2005). To overcome this problem, two shorter overlapping constructs were used for further NMR analysis: KKT4^115–174^, the minimal microtubule-binding domain which retains reduced microtubule-binding activities (Llauró et al., 2018), and KKT4^145–232^, which was identified as a stable fragment in trypsin digests of KKT4^115–343^ (Ludzia et al., 2020). The 2D ^1^H-^15^N correlation spectra of these constructs contained peaks for essentially all residues (Ludzia et al., 2020). In addition, the superposition of the NMR spectra of KKT4^115–174^ and KKT4^145–232^ with that of KKT4^115–232^ showed good agreement in peak positions (Supplemental Figure 3A, B). This indicates that the two shorter constructs retain the structural and dynamical properties observed in the longer fragment, thereby justifying our dissection approach. The chemical shifts of KKT4^115–174^ and KKT4^145–232^ were analysed using TALOS-N (Shen and Bax, 2013). Both of these fragments revealed significant amounts of secondary structure (Figure 2C, D). For KKT4^145–232^, a continuous helix was observed from 152 to 225 (Figure 2C). For KKT4^115–174^, two helices, encompassing residues 121 to 139 and 152 to 172, separated by an unstructured linker were observed (Figure 2D).

The presence of an unstructured linker in KKT4^115–174^ suggests flexibility between the two helices. To confirm this, we used the {^1^H}–^15^N heteronuclear NOE to probe backbone dynamics in KKT4^115–174^ and KKT4^145–232^ (Figure 2E). The {^1^H}–^15^N NOE ratios for KKT4^115–174^ and KKT4^145–232^ are consistent with the predicted secondary structure. {^1^H}–^15^N NOE ratios of greater than ∼0.6, characteristic of a structured backbone, were found for most residues that were predicted to be helical (Figure 2C, D). In KKT4^115–174^, residues 140–151, which were not predicted to be helical, had lower {^1^H}–^15^N NOE ratios, confirming the presence of a disordered dynamic linker between the two helices (Figure 2E). Interestingly, higher {^1^H}–^15^N NOE ratios were observed at the start of the 2^nd^ helix in KKT4^115–174^ than for this region in KKT4^145–232^. This may indicate that the presence of the full linker and/or the 1^st^ helix contributes to the stability of the 2^nd^ helix. The spectrum of KKT4^115–343^ contained peaks for residues 115–150, which correspond to the 1^st^ helix and part of the flexible linker in KKT4^115–174^ (Supplemental Figure 2B). The {^1^H}–^15^N NOE ratios for these residues in the KKT4^115–343^ construct agree with the values measured for KKT4^115–174^, indicating that this region has the same dynamic properties in the two constructs. In summary, the microtubule-binding domain of KKT4 is composed of two helices, encompassing residues 121 to 139 and 152 to 225, separated by a 12-residue flexible linker, followed by a ∼120-residue positively charged disordered region (predicted isoelectric point for residues 226–343 is 10.1).

### Crystal structures of *T. cruzi* KKT4^117–218^ and *L. mexicana* KKT4^184–284^

To characterise the structure of the microtubule-binding domain in greater detail, we designed additional constructs for crystallography based on the regions shown to be helical by NMR. Although we still failed to obtain diffraction-quality crystals from *T. brucei*, a crystal structure of KKT4^117–218^ from *T. cruzi* was solved at 1.9 Å resolution (Table 2). *Tc*KKT4^117–218^ is homologous to KKT4^120–224^ in *T. brucei* (Figure 3A, Supplemental Figure 5), which has the two helical regions identified by NMR. The structure consists of two ∼150 Å long parallel *α*-helices organised in a left-handed coiled-coil dimer (Figure 3B); the helical structure starts at L118 and ends at D215. The coiled coil consists of 8 regular heptad repeats starting at Y121 and ending at K176. Analysis with TWISTER (Strelkov and Burkhard, 2002) shows that the six central heptads are characterised by an inter-helical distance of 4.84 ± 0.06 Å and a coiled-coil pitch (the periodicity of the coiled coil) of 136 ± 26 Å; the latter value is close to the theoretical value of 135 Å for a coiled coil. If the 1^st^ and 8^th^ heptad are included, the pitch increases to 163 ± 60 Å indicating less supercoiling in these terminal heptads. After K176, the helices move apart (inter-helical distance increases to 5.66 ± 0.15 Å) and the coiled-coil pitch increases significantly, indicating a loss of supercoiling.

**Figure 3.**
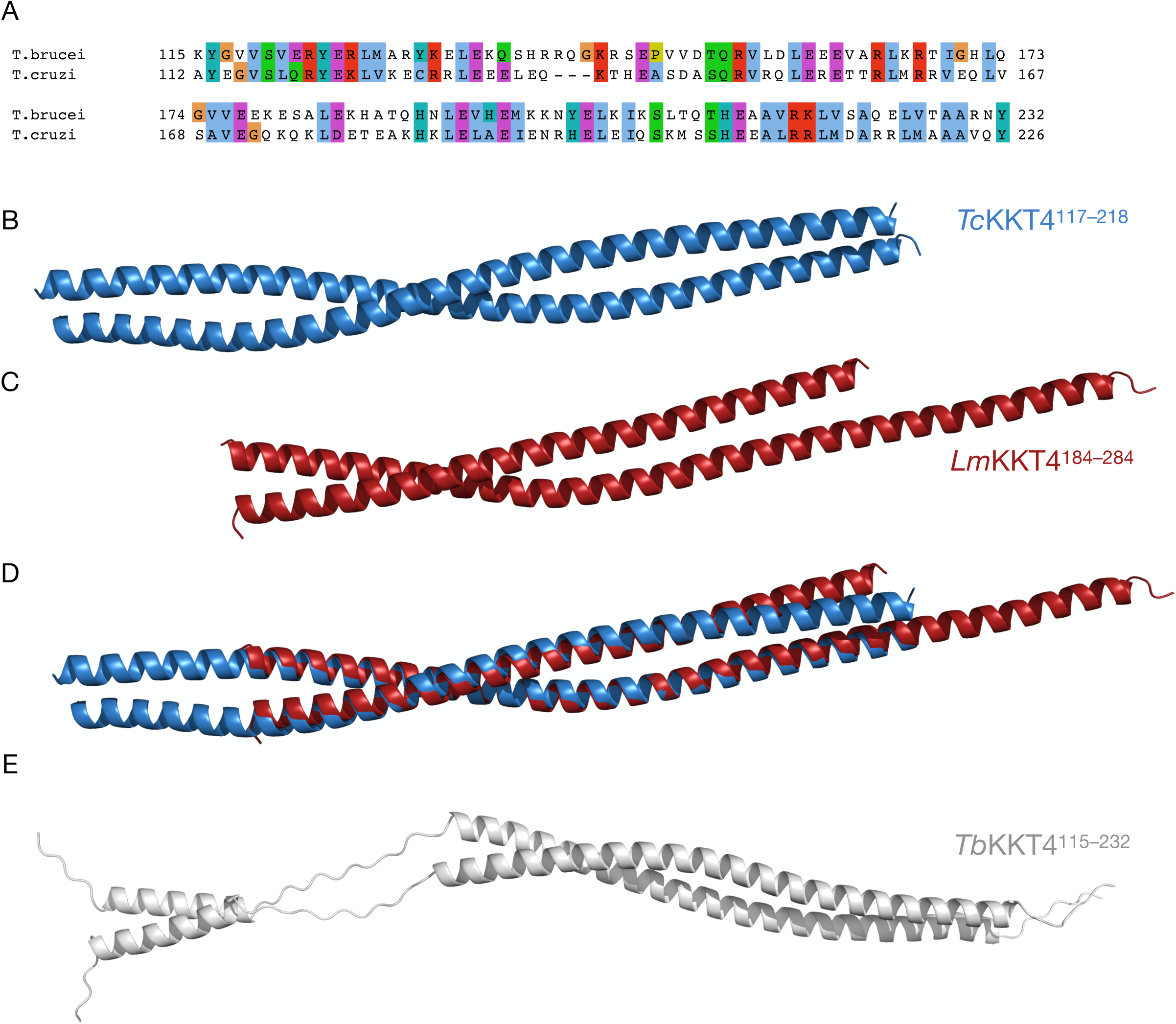
Crystal structures of *T. cruzi* KKT4^117–218^ and *L. mexicana* KKT4^184–284^ reveal coiled coils. (A) Sequence alignment of the KKT4 microtubule-binding domain between *Trypanosoma brucei* and *Trypanosoma cruzi* (Sylvio X10). (B) Ribbon model of the *T. cruzi* (Sylvio X10) KKT4^117–218^ backbone. The N-terminus of the structure starts from the left. (C)Ribbon model of the *Leishmania mexicana* KKT4^184–284^ backbone. The N-terminus of the structure starts from the left. (D) Superposition of the *T. cruzi* KKT4^117–218^ (blue) and *L. mexicana* KKT4^184–284^ (ruby) structures. An RMSD (1.03 Å) was calculated using the *align* function in Pymol (DeLano, 2002). (E) Ribbon representation of the *Trypanosoma brucei* KKT4^115–232^ structural model generated using MODELLER (Webb and Sali, 2016). The model is based on NMR data (see Figure 2) showing two coiled-coil segments separated by an unstructured linker and flanked by small disordered regions at the N-and C-termini of the molecule.

**Table 2.** Data collection, refinement statistics

|  | *T. cruzi* KKT4<sup>117–218</sup> | *L. mexicana* KKT4<sup>184–284</sup> | *T. brucei* KKT4<sup>463–645</sup> |
| --- | --- | --- | --- |
| **Data collection** |  |  |  |
| Beamline | Diamond Light Source I03 | Diamond Light Source I24 | Diamond Light Source I24 |
| Wavelength (Å) | 0.9760 | 0.91587 | 0.96861 |
| Resolution range (Å) | 67.96–1.90 | 61.15–1.90 | 45.60–1.57 |
| Space group (Z) | *P* 1 2<sub>1</sub> 1 | *P* 1 2<sub>1</sub> 1 | *P* 2<sub>1</sub> 2<sub>1</sub> 2<sub>1</sub> |
| Unit cell | 33.62 Å 25.31 Å 136.88 Å<br>90° 96.81° 90° | 31.31 Å 37.71 Å 122.39 Å<br>90° 92.17° 90° | 46.37 Å 61.63 Å 67.78 Å<br>90° 90° 90° |
| Unique reflections | 16812 | 22522 | 27032 |
| Multiplicity | 6.1 | 6.5 | 10.0 |
| Completeness (%) | 90.4 | 98.3 | 96.8 |
| *I*/σ*I* | 4.28 | 2.84 | 2.86 |
| Wilson B-factor (Å<sup>2</sup>) | 24.4 | 17.6 | 15.7 |
| CC<sub>1/2</sub> | 0.998 | 0.997 | 0.997 |
| **Refinement** |  |  |  |
| No. reflections | 16788 | 22502 | 26945 |
| R<sub>work</sub> | 0.241 | 0.199 | 0.173 |
| R<sub>free</sub> | 0.258 | 0.232 | 0.192 |
| Number of atoms | 1871 | 1816 | 1494 |
| Protein | 1670 | 1488 | 1270 |
| Solvent | 201 | 328 | 224 |
| RMS bonds (Å) | 0.006 | 0.014 | 0.011 |
| RMS angles (°) | 0.97 | 1.45 | 1.46 |
| Ramachandran favoured (%) | 100.00 | 100.00 | 98.68 |
| Ramachandran allowed( %) | 0.00 | 0.00 | 1.32 |
| Ramachandran outliers (%) | 0.00 | 0.00 | 0.00 |
| Average B-factor (Å<sup>2</sup>) | 36.0 | 30.0 | 22.0 |

To test structural conservation in other kinetoplastid species, we attempted to solve the crystal structure of the KKT4 coiled-coil fragment from *Leishmania mexicana*. We were unable to obtain diffraction-quality crystals of fragments that had the N-terminal residues of the microtubule-binding domain (115–140 in *T. brucei*). Instead, we crystallised and solved a 1.9 Å structure of *Lm*KKT4^184–284^ (Table 2), which corresponds to residues 141–244 in *T. brucei* (Supplemental Figure 5). We note that the expressed protein contained an additional 23 residues from the expression vector at its C-terminus due to a cloning error (see Materials and Methods). We failed to obtain crystals for the construct that ends at Q284. Similarly to *Tc*KKT4^117–218^, *Lm*KKT4^184–284^ has helices arranged in a parallel coiled-coil fold (Figure 3C). *Lm*KKT4^184–284^ has 6 regular heptad repeats from the N-terminus to Q224, and after that point the helices move apart and lose their supercoiling as observed for *Tc*KKT4^117–218^. The two chains in *Lm*KKT4^184–284^ differ in length with helices spanning T184 to A289 in one chain and T184 to R257 in the other. The region beyond I245, where the helices have moved away from each other, had high B-factors in both chains suggesting enhanced flexibility (Supplemental Figure 6). The electron density of the shorter helix disappears at Q258, most likely due to the disordered nature of the protein backbone in that region. Interestingly, the end of the longer helix, which contains extra residues from the expression vector, makes seemingly stabilising contacts with other molecules in the crystal lattice, which potentially explains why our attempts to obtain diffraction-quality crystals for the proper construct ending at Q284 failed.

A DALI search (Holm, 2019) of the protein data bank (PDB) for structural homologues of *Tc*KKT4^117–218^ and *Lm*KKT4^184–284^ identified similarity to a number of coiled-coil proteins (Supplemental Tables 1 and 2). Superposition of the coiled coils from *T. cruzi* and *L. mexicana* revealed a good structural match with an RMSD of 1.03 Å (for 123 C*α*) (Figure 3D), confirming the conservation of the coiled-coil structure in these species. The helical regions in the crystal structures of *Tc*KKT4^117–218^ and *Lm*KKT4^184–284^ match those in *T. brucei* KKT4 in solution identified by NMR. However, in both crystal structures, we did not find a flexible linker within the coiled coils that we identified in *T. brucei* KKT4 using NMR. Instead, we observed high B-factors (Supplemental Figure 6) for the region where we expect to find an unstructured linker in the *Tc*KKT4 structure based on the sequence alignment (Supplemental Figure 5). We speculate that the lack of a flexible linker in *T. cruzi* and *L. mexicana* crystal structures may be either due to the stabilising contacts within the crystal lattice or structural differences of KKT4 between *T. brucei* and the other two kinetoplastids.

### Modelling of the *T. brucei* KKT4 coiled coil

Although we could not solve a crystal structure for *T. brucei* KKT4, our NMR analysis identified helical structure in the N-terminal part of the microtubule-binding domain. Together with the finding that *T. cruzi* and *L. mexicana* homologues have dimeric coiled-coil structures, we next aimed to determine if the helices in *T. brucei* KKT4 adopt similar structures. Although NMR chemical shifts alone cannot distinguish between undistorted and supercoiled helices, residual dipolar couplings (RDCs) measured for partially aligned protein samples can provide this information. RDCs are very sensitive to N-H bond vector orientation and are suited to characterising the bending of *α*-helices that occurs in coiled-coil structures. In a supercoiled helix, the helical turns at the packing interface (residues *a, d, e, g* in the heptad repeat) are slightly compressed while those facing outside (*b, c, f*) are stretched, which leads to a periodic variation in the RDCs within the heptad repeat (Schnell et al., 2005).

RDCs were measured for KKT4^115–174^ partially aligned in 3% PEG-hexanol (Figure 4). Both helices in KKT4^115–174^ show large positive RDCs while the linker region and the N-and C-terminal residues show RDCs close to 0. The RDCs for both helices show periodic variation that is consistent with a coiled-coil structure. The flexible linker separating the helices is dynamic, sampling multiple conformations, and this leads to averaging of its RDCs to values close to 0. The different values of the RDCs for the two helices likely reflect the presence of the flexible linker, which means that the two helices can move independently and the molecule does not align as a single rigid unit.

**Figure 4.**
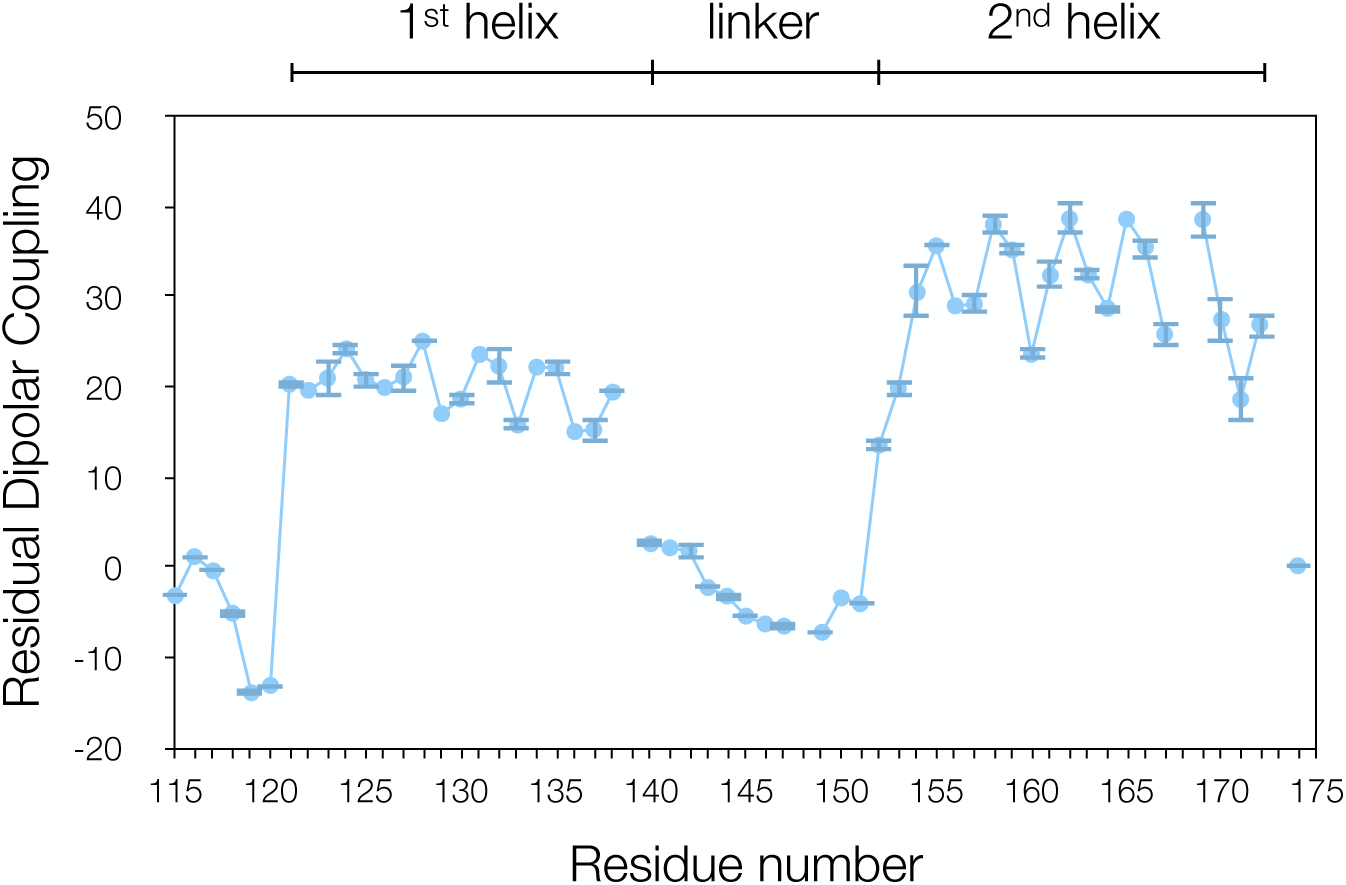
Residual dipolar couplings of *T. brucei* KKT4^115–174^ reveal coiled coils. Experimental ^1^H-^15^N residual dipolar couplings (RDCs) for KKT4^115–174^, measured in 3% C12E6/n-hexanol, are plotted as a function of sequence. The RDCs for both helices show periodic variation that is consistent with a coiled-coil structure. The RDCs close to 0 for the linker region indicate that this region is dynamic.

Using the *T. cruzi* X-ray structure as a model, we tested different ways of fitting the helices identified by NMR into a coiled-coil structure by optimising the fit between RDCs predicted from the X-ray structure and the experimental RDCs (Supplemental Figure 7A, B). For both helices in *T. brucei*, good fits were found when they were placed at several positions within the first half of the *T. cruzi* sequence, corresponding to the regular coiled-coil structure found between residues 121 and 176. Poorer agreement was obtained when the *T. brucei* helices were placed within the C-terminal half of the *T. cruzi* structure that showed less supercoiling. For the 2^nd^ helix, the fit of the experimental RDCs suggests an offset of −6 residues between the *Tb* and *Tc* sequences (Supplemental Figure 7B), in agreement with the sequence alignment of these two proteins (Figure 3A); this places hydrophobic residues in *T. brucei* in positions *a/d* of the heptad repeat in the *T. cruzi* structure. For the 1^st^ helix, two solutions, with sequence offsets of −3 or 0 residues, are found from the fitting of experimental RDCs (Supplemental Figure 7A). The sequence offset of 0 was excluded because this would place L127 and L134 in heptad position *g* away from the helix packing interface and would not give rise to the experimentally observed upfield shifts for the methyl groups that arise from packing with Y124 and Y131 at the interface (Supplemental Figure 7C, D). The sequence offset of −3 to be determined from the RDC and chemical shift data is in agreement with the alignment of the *Tb* and *Tc* sequences (Figure 3A) in the region of the 1^st^ helix.

A homology model for the two coiled-coil regions of *T. brucei* KKT4^115–232^ was built using the *T. cruzi* coiled-coil X-ray structure, and the sequence alignments derived from the RDC data (Webb and Sali, 2016). Random extended structures for the flexible N- and C-termini and for the inter-helix linker, all identified as flexible on the basis of the heteronuclear NOE data, were generated; these represent possible conformations that these residues might sample in solution. These coordinates were merged with the coiled-coil homology models to generate an overall model for *T. brucei* KKT4^115–232^ (Figure 3E). This model allows the properties of the surface of *T. brucei* KKT4^115–232^ to be investigated.

### Mapping the microtubule-binding interface of KKT4

Microtubule interaction is often mediated by the electrostatic effects of surface charges (Ciferri et al., 2008). In fact, we previously showed that a charge-reversal mutant that replaced three basic residues with acidic residues (R123E, K132E, R154E) severely reduced the microtubule-binding activity of *T. brucei* KKT4^115–343^ (Llauró et al., 2018). To understand the electrostatic surface of the coiled coil in *T. brucei* KKT4, we used the homology model described above and calculated the electrostatic surface potential using the APBS electrostatic plugin in Pymol (Jurrus et al., 2018). The structural model of *T. brucei* KKT4^115–232^ revealed positively charged regions in the N-terminus of the coiled-coil structure (Supplemental Figure 8A), with basic residues exposed on the protein surface (e.g. R123, R126, R130, K132, K136, R140, K199, K204, K206, R217 and K218) (Supplemental Figure 8B). To test the importance of positively charged residues for the microtubule binding, we systematically generated single mutants of KKT4^115–343^ where lysine and arginine residues that are present within the N-terminal region (115–232) of the microtubule-binding domain were replaced with glutamic or aspartic acid. Microtubule binding of these mutants was compared to that of wild-type KKT4^115–343^ using co-sedimentation assays. We found that mutating any of residues R123, R126, R130, K136, R140, R141, K144 and R145 significantly reduced the microtubule-binding affinity (Figure 5, Supplemental Figure 8B). These residues are located within KKT4^115–174^, which was previously identified as the minimal microtubule-binding domain that had weaker microtubule-binding activities compared to KKT4^115–343^ (Llauró et al., 2018). In contrast, mutating basic residues located in the second, longer helix in the coiled-coil of the protein (K154, R164, K166, R167, K179, K198, K199, K204, K206, R217, K218, and R230) had only mild effects on the microtubule-binding activities (Figure 5). To test if mutations affected the stability of the structure, 1D ^1^H NMR spectra were collected for mutants that had reduced affinity to microtubules (R126, R130, K136, R140, R141, K144); these confirmed that the mutations did not alter the protein structure (data not shown). Together with our previous analysis (Llauró et al., 2018), these results confirmed the importance of positively charged residues for microtubule-binding activities and revealed that the primary microtubule-binding interface of KKT4 is likely located in the N-terminus of the coiled-coil.

**Figure 5.**
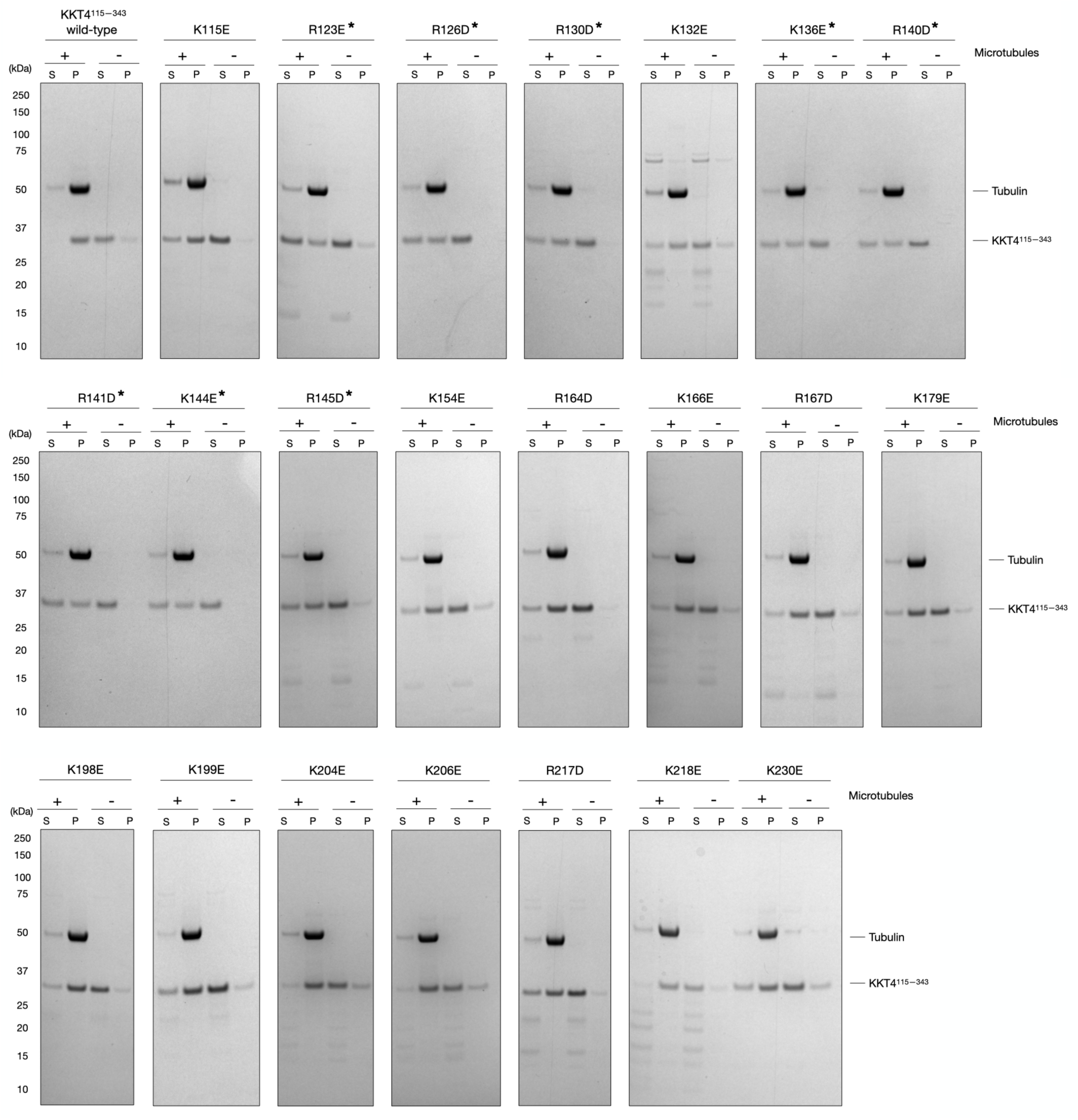
Mapping the microtubule-binding interface of KKT4. Microtubule co-sedimentation assay of KKT4^115–343^ charge-reversal mutants, showing that mutations in its N-terminal region reduce the microtubule-binding affinity. The intensity differences of KKT4 bands between supernatant and pellet fractions were calculated using ImageJ (Schneider et al., 2012); the mutants that show ∼50% reduction in the microtubule-binding are indicated with an asterisk (*). S and P correspond to supernatant and pellet fractions, respectively.

### KKT4 has DNA-binding activities

In many species, kinetochore assembly is regulated during the cell cycle. In humans, the constitutive centromere-associated network (CCAN) is composed of 16 chromatin-proximal kinetochore proteins that act as a platform for kinetochore assembly by interacting with CENP-A nucleosomes (Cheeseman and Desai, 2008). Most of the CCAN components constitutively localise at kinetochores and some of them have DNA-binding activities. In contrast, microtubule-binding kinetochore components localise only during mitosis. In *T. brucei* kinetochore assembly is regulated during the cell cycle. However, the microtubule-binding protein KKT4 localises at kinetochores in a constitutive manner (Akiyoshi and Gull, 2014). Interestingly, the C-terminus of KKT4 is predicted to be a tandem BRCT domain (Akiyoshi and Gull, 2014). BRCT domains are found in a number of prokaryotic and eukaryotic proteins with various biological functions including DNA or RNA binding (Zhang et al., 1998, Yu et al., 2003, Leung and Glover, 2011). We previously observed significant DNA contamination during the purification of full-length KKT4 from insect cells (Llauró et al., 2018), suggesting that KKT4 might have DNA-binding activity, possibly via its BRCT domain. To test this, we employed a fluorescence anisotropy assay (Rossi and Taylor, 2011) using a 50-bp double-stranded DNA, labelled at the 5’ end with 6-carboxyfluorescein (6-FAM). We found that the full-length KKT4 (KKT4^2–645^) strongly bound DNA (KD = 11 nM) (Figure 6), while the KKT4 BRCT domain (KKT4^463–645^) failed to saturate the DNA signal under the same conditions used for the full-length protein (Figure 6). These results suggest that the BRCT domain of KKT4 does not bind DNA tightly and that the high-affinity DNA-binding domain is located elsewhere in KKT4.

**Figure 6.**
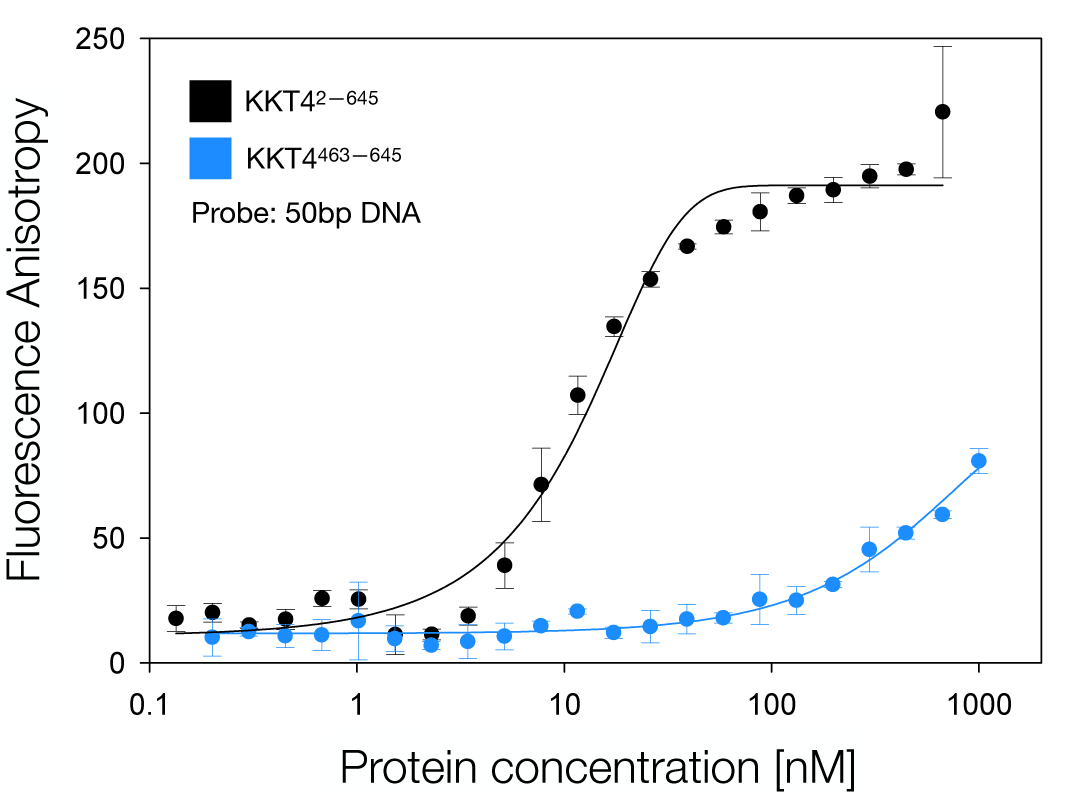
KKT4 has DNA-binding activities. Measured anisotropy is plotted against KKT4^2–645^ and KKT4^463–645^ protein concentration in the fluorescence anisotropy assay using a 50-bp DNA probe. The DNA sequence (∼50% GC content) used in this assay is part of the centromeric sequence (CIR147) in *T. brucei*. The KD was calculated using non-linear regression using SigmaPlot (Monks, 2002); the fit is shown as a solid line.

### Crystal structure of the *T. brucei* KKT4 BRCT domain

To understand the function of the KKT4 BRCT domain, we used X-ray crystallography to solve its structure. KKT4^463–645^ yielded crystals that diffracted to a resolution of 1.6 Å (Table 2). BRCT domains typically comprise between 90 and 100 amino acids with the *βαββαβα* secondary structure topology (Leung and Glover, 2011). The structure of KKT4^463–645^ revealed tandem BRCT domains (Figure 7A). The N-terminal full domain (BRCT1) consists of a central 4-stranded β-sheet flanked by 2 *α*-helices on one side of the sheet and one *α*-helix on the opposite side. The smaller domain (BRCT2) consists of 2 *α*-helices and 3 β-strands, missing a β-strand and an *α*-helix in the C-terminus (Figure 7A). No electron density for residues 463–473, 519–523 and 617–625 was observed, suggesting that these regions are flexible. A search for structural homologues using DALI (Holm, 2019) revealed similarity to several BRCT-containing proteins with Breast cancer associated protein 1 (BRCA1) as one of the top hits (Supplemental Table 3). The tandem BRCT domains in BRCA1 have a highly conserved pocket that binds phosphopeptides (Clapperton et al., 2004, Shiozaki et al., 2004, Williams et al., 2004). Superposition of KKT4 BRCT1 with the N-terminal BRCT domain of *H. sapiens* BRCA1 showed a good structural match with an RMSD of 1.11 Å (65 to 65 C*α*) (Figure 7B), suggesting that the KKT4 BRCT domain likely binds phosphopeptides. In fact, in the structure of the KKT4 BRCT domain, we observed additional electron density, likely arising from a sulphate ion that may mimic a bound phosphate group. The sulphate ion is coordinated by three residues in the pocket, T494, S495 and K543 (Figure 7C), which correspond to the key amino acids known to interact with phosphopeptides in other BRCT domains (e.g. S1655, G1656, and K1702 in human BRCA1) (Williams et al., 2004). These results suggest that the BRCT domain of KKT4 likely functions as a phosphorylation-dependent protein-protein interaction domain rather than a DNA-binding domain.

**Figure 7.**
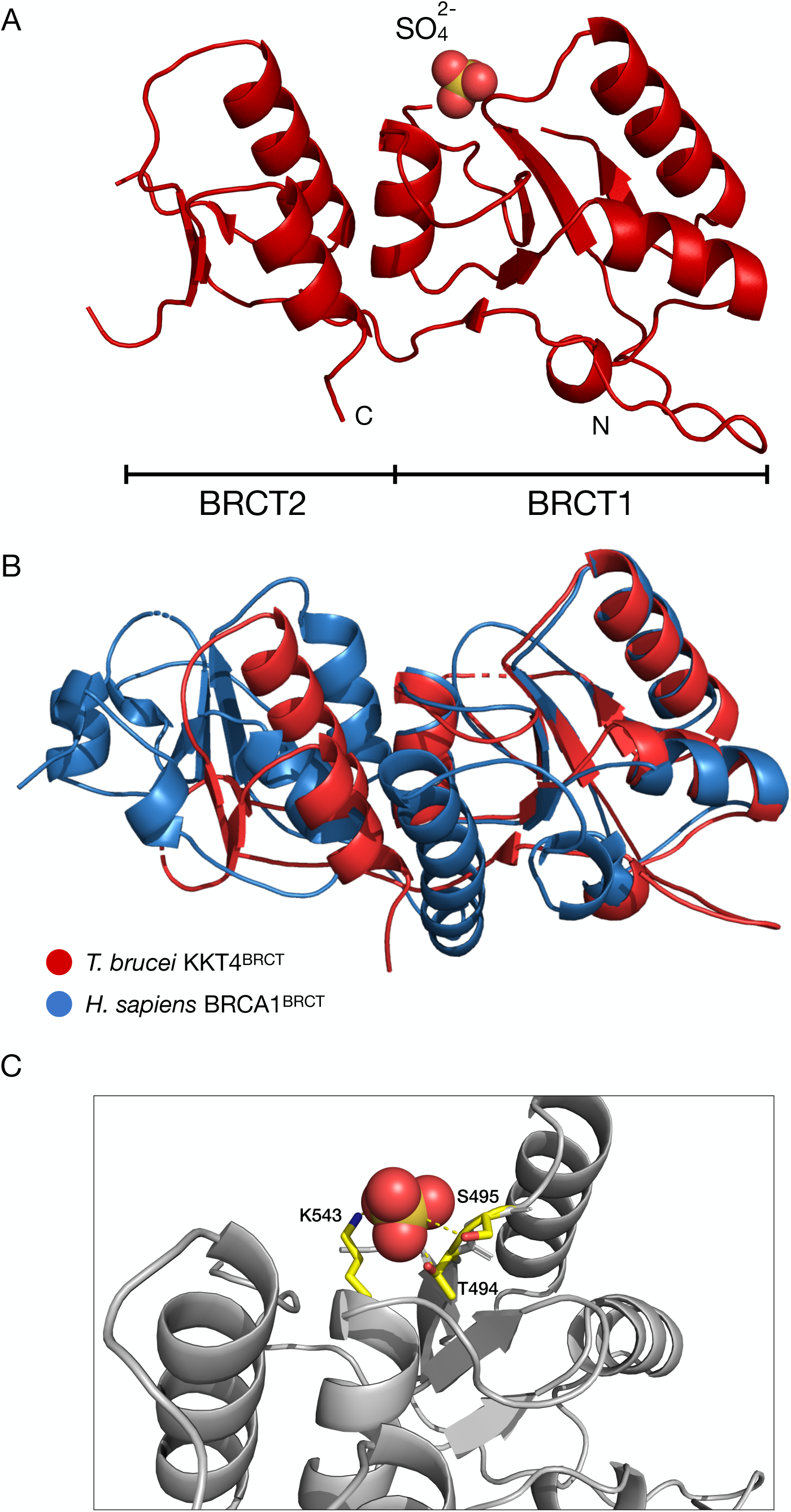
Crystal structure of the tandem BRCT domains of KKT4 reveals a putative phosphopeptide-binding pocket. (A) Ribbon representation of KKT4 BRCT domain (KKT4^463–645^). The N- and C-terminus are indicated by N and C respectively. The residues for which the electron density was not visible are not shown in the figure. (B) Superposition of the KKT4 BRCT domain (red, PDB: 6ZPK) with the BRCA1 BRCT domain (blue, PDB: 3FA2), highlighting the absence of a β-strand and an α-helix in the C-terminus of KKT4 BRCT. The RMSD (1.11 Å) was calculated using the *super* function in Pymol (DeLano, 2002). (C)Close-up view showing coordination of a sulphate ion by T494, S495 and K543 (side chains of these residues are shown as yellow sticks).

### The KKT4 BRCT domain interacts with the microtubule-binding region

To obtain further structural information on KKT4, we used crosslinking mass spectrometry (XL-MS), which can identify interaction surfaces between partner proteins or within the same molecule (Mattson et al., 1993, Leitner et al., 2016). We first carried out crosslinking on full-length KKT4 using bis(sulfosuccinimidyl)suberate (BS^3^), a homo-bifunctional crosslinker that reacts with primary amines. BS^3^ covalently links pairs of lysines that are within 26–30 Å on the protein surface. XL-MS of KKT4 resulted in numerous crosslinks across the molecule (Figure 8A, top). We then used a different type of crosslinker, zero-length 1-ethyl-3-(3-dimethylaminopropyl)carbodiimide hydrochloride (EDC) and *N*-Hydroxysulfosuccinimide sodium salt (Sulfo-NHS) that activates carboxyl groups for reaction with primary amines (Figure 8A, bottom). The pattern of crosslinks was similar to that obtained with BS^3^. Interestingly, a number of crosslinks were identified between the BRCT domain and the microtubule-binding domain using both BS^3^ (K115/K499, K115/K521, K115/K543, K132/K618, K199/K521, K206/K521, K218/K499, K218/K510, K218/K521, K218/K618) and EDC/Sulfo-NHS (K115/D645, K132/E575, K132/D645, E178/K521, K204/D645, K206/E573, K206/D645 K218/E573, K218/D645). These results suggested that the KKT4 BRCT domain interacts directly with the microtubule-binding domain.

**Figure 8.**
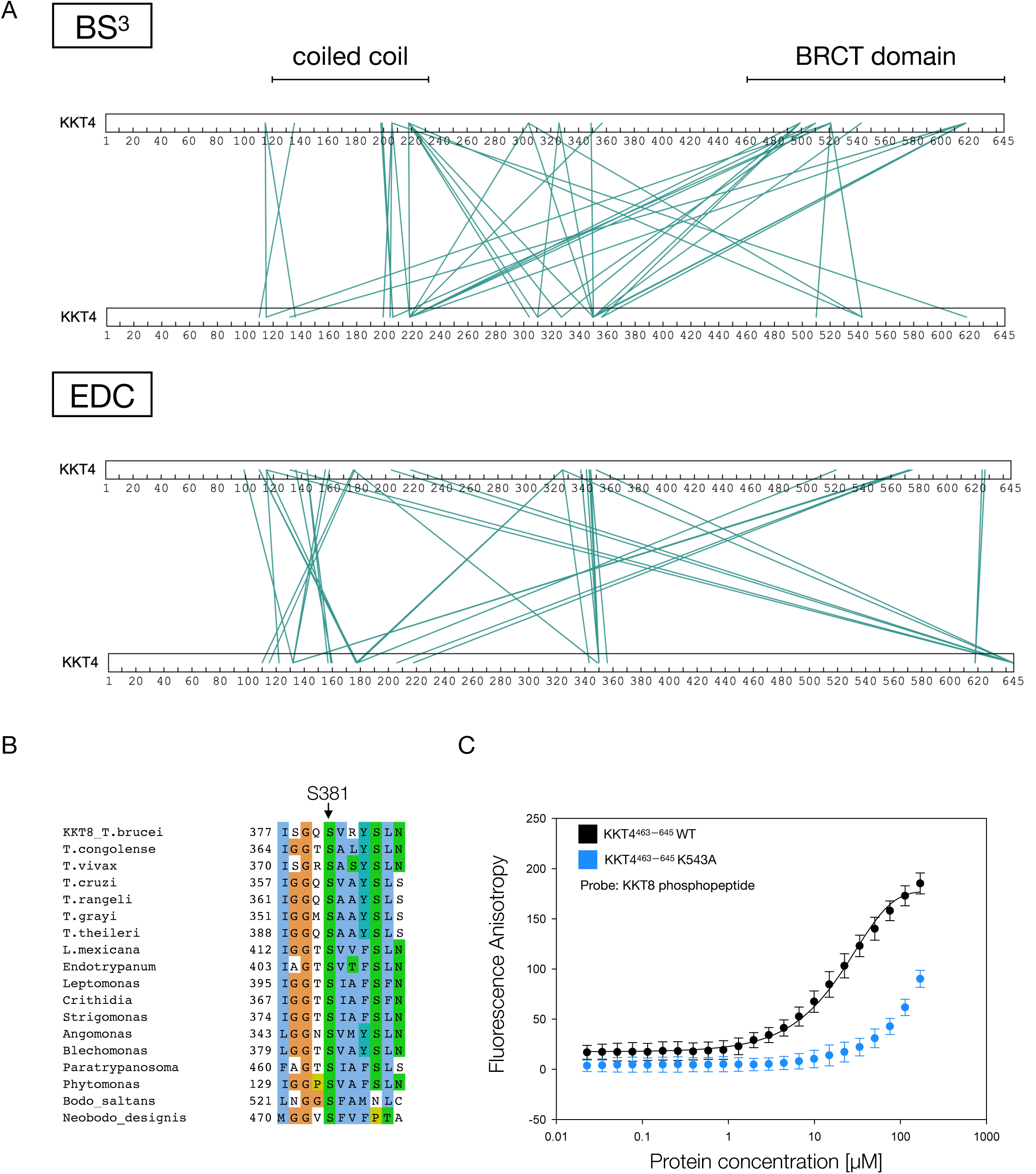
The KKT4 BRCT domain interacts with the microtubule-binding domain and a phosphopeptide derived from KKT8. (A) Crosslinking mass spectrometry of full-length KKT4 using BS^3^ and EDC/Sulfo-NHS. The green lines indicate pairs of crosslinked residues. For the purposes of clarity, two molecules of KKT4 are shown. Note that crosslinks between the top and bottom KKT4 do not necessarily mean that crosslinks formed in between two separate molecules because it was not possible to distinguish between inter- and intra-molecule crosslinks in this experiment (except for those inter-molecule crosslinks formed between the same residues). A complete list of identified crosslinks is shown in Supplemental Table 4 and 5. (B) Multiple sequence alignment of KKT8, showing the conservation of *T. brucei* S381. KKT8 protein sequences from various kinetoplastids were aligned using MAFFT (Katoh et al., 2019) and visualised with the CLUSTALX colouring scheme in Jalview (Waterhouse et al., 2009). (C)Fluorescence anisotropy assay showing the KKT4 BRCT domain binding to a KKT8 phosphopeptide (wild type in black and K543A mutant in blue). The KD (30 µM) for the wild type BRCT domain was calculated using non-linear regression using SigmaPlot (Monks, 2002); the fit is shown as a solid line.

To confirm this, we collected 2D ^1^H-^15^N BEST TROSY spectra of ^15^N-KKT4^115–232^ mixed with the unlabelled KKT4 BRCT domain (residues 463–645, KKT4^BRCT^) (Supplemental Figure 9A, B). In these experiments, the peaks corresponding to residues from KKT4^115–232^ that interact with the BRCT domain should show chemical shift changes due to the altered environment, while those corresponding to residues that are not involved in binding remain unaffected. Comparison of the KKT4^115–232^ spectrum with and without KKT4^BRCT^ revealed several residue-specific changes; observed shifts are plotted as a function of sequence in Supplemental Figure 9A. The peaks that showed the largest shifts correspond to K115, Y116, G117, V118, V119, S120, V121, E122 and R123 (Supplemental Figure 9). It is interesting that one of the residues that shifted, K115, was shown to crosslink with K543 from the BRCT domain. We repeated the experiment with a shorter KKT4 construct, KKT4^115–174^, and obtained similar results (Supplemental Figure 9A). Although crosslinks were also observed between the BRCT domain and residues E178, K199, K204, K206 and K218, these residues did not give observable peaks in the spectra of KKT4^115–232^ so we could not confirm the interaction by NMR. When similar experiments were performed with ^15^N-KKT4^145–232^ and KKT4^BRCT^, no significant changes in chemical shift were observed (Supplemental Figure 9A). These results suggest that the interaction between the BRCT domain and the microtubule-binding domain of KKT4 depends on residues at the N-terminus of the microtubule-binding domain.

### The KKT4 BRCT domain is a phosphopeptide-binding domain

The interaction identified above between the N-terminal region of the microtubule-binding domain and the BRCT domain appears to be relatively weak based on the NMR data. Although the interacting N-terminal region contains a serine residue (S120 with R123 at +3 position), it does not match a typical consensus motif (pS/pT)-x-x-(F/Y/I/L) for BRCT binding (Manke et al., 2003, Yu et al., 2003) and is unlikely to bind the phosphopeptide-binding pocket. To identify other potential binding partners for the KKT4 BRCT domain, sequences of kinetochore proteins were searched for the consensus motif. Among those proteins that co-purified with KKT4 (Akiyoshi and Gull, 2014), we identified possible motifs in KKT7 (SVTF, residues 65–68), KKT8 (SVRY, residues 381–384), and KKT12 (SILL, residues 192–195), which are highly conserved among kinetoplastids (Figure 8B and data not shown). Fluorescently-labelled phosphopeptides derived from these proteins were tested for KKT4^BRCT^ binding using a fluorescence anisotropy assay. The peptide derived from KKT8 bound KKT4^BRCT^ with a KD of ∼30 µM (Figure 8B). The other two peptides failed to bind KKT4^BRCT^ with a similar affinity (data not shown). Importantly, replacement of K543 that is located in the putative phosphopeptide-binding site in KKT4^BRCT^ with alanine decreased the binding affinity by roughly an order of magnitude (Figure 8C). These results show that KKT4 BRCT is a phosphopeptide-binding domain and identify KKT8 as a potential interaction partner of KKT4.

## Discussion

Many kinetochore-localised microtubule-binding proteins have been characterised in other model organisms, such as the Ndc80, Ska, and Dam1 complexes, SKAP/Astrin, CENP-E, CENP-F, MCAK, INCENP, XMAP215, and dyneins (Maiato et al., 2004, Foley and Kapoor, 2013, Musacchio and Desai, 2017). Besides folded domains, many microtubule-binding proteins have predicted disordered regions that enhance their binding affinity (Guimaraes et al., 2008, Friese et al., 2016, Volkov, 2020). It is noteworthy that the predicted disorder has not been confirmed in most cases. Similarly, KKT4 has a predicted disordered segment, which is not sufficient to bind microtubules but enhances the binding affinity (Llauró et al., 2018). In this study, we used NMR to confirm that this region is indeed disordered (Figure 9). We also found that the N-terminal half of the KKT4 microtubule-binding domain has an elongated coiled-coil fold (Figure 9). Although a number of kinetochore proteins have coiled coils, microtubule-binding domains are typically located elsewhere, such as the calponin-homology domain (CH domain) for Ndc80/Nuf2 (Wei et al., 2007, Ciferri et al., 2008). In the case of SKAP/Astrin, which also has predicted coiled coils, it has been shown that the coiled-coil segment is unable to interact with microtubules on its own and that microtubule binding requires the N-terminal disordered fragment (Friese et al., 2016). Our mutagenesis analysis suggested that KKT4 binds microtubules through the N-terminal region of the coiled coil. In the future, it will be important to directly visualise the microtubule-binding interface using methods such as electron microscopy.

**Figure 9.**
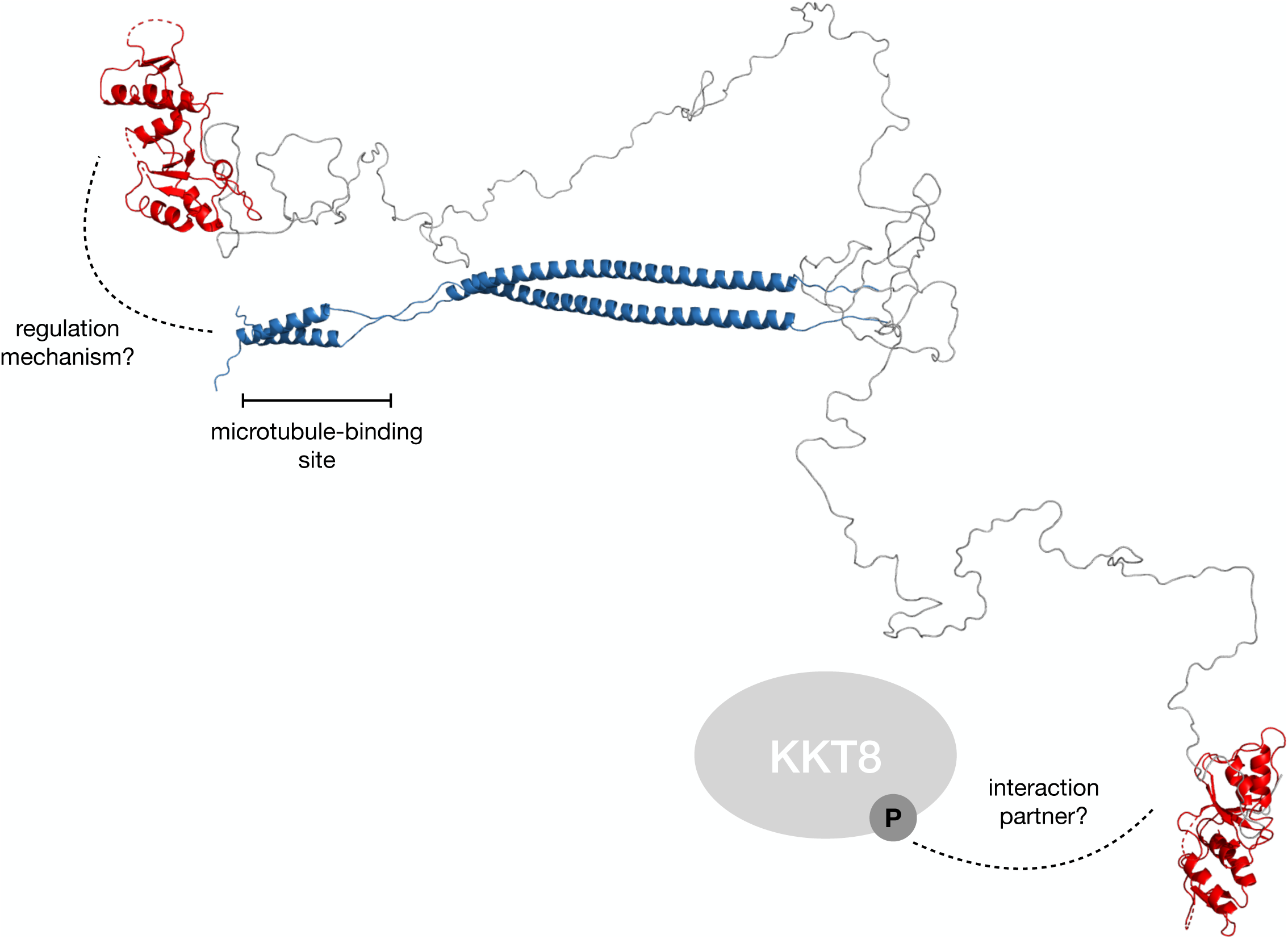
Structural and functional model of *T. brucei* KKT4^115–645^. This model was generated using X-ray crystallography and NMR data. The residues that are important for the KKT4 microtubule-binding activity are located in the region indicated by the bar. The model shows two possible orientations of the BRCT domain (shown in red) with respect to the coiled coil (shown in blue) based on random conformations for the disordered region between Q233 and T473. These disordered conformations allow the BRCT domain to interact with the N-terminal region of the coiled coil. However, alternative conformations in which the BRCT domain is distant from the coiled coil are also possible, which might favour interactions with other kinetochore proteins such as phosphorylated KKT8 (shown as a grey ellipsoid).

It remains unknown whether (and how) microtubule-binding activities of KKT4 are regulated. Interestingly, we found that the KKT4 BRCT domain interacts with the N-terminal part of the microtubule-binding domain (Figure 9), raising a possibility that this interaction might affect the microtubule-binding activities of KKT4. Alternatively, the observed interaction might regulate other activities of KKT4. We previously showed that KKT4 co-purifies with the APC/C subunits (Akiyoshi and Gull, 2014). Furthermore, although trypanosomes lack a canonical spindle checkpoint (Ploubidou et al., 1999), cells could control the cell cycle progression by regulating the level of cyclin B in the nucleus (Hayashi and Akiyoshi, 2018). We therefore speculate that the interaction between the BRCT domain and the microtubule-binding domain might be governed by the attachment status, which in turn controls APC/C activities and the cell cycle progression.

In other eukaryotes, Aurora B kinase plays an important role in regulating kinetochore-microtubule attachment by phosphorylating microtubule-binding kinetochore proteins, including Ndc80 and the Ska complex (Cheeseman et al., 2002, Tien et al., 2010, Chan et al., 2012, Redli et al., 2016). Although Aurora B is conserved in kinetoplastids, it remains unclear whether it regulates kinetochore-microtubule attachments (Tu et al., 2006). Our preliminary *in vitro* kinase assay failed to find evidence that *T. brucei* Aurora B phosphorylates KKT4 (data not shown). In contrast, we previously showed that KKT4 is phosphorylated by the KKT10 kinase, which localises at kinetochores until the onset of anaphase and promotes the metaphase-to-anaphase transition (Ishii and Akiyoshi, 2020). A phospho-deficient KKT4^S477A^ mutant failed to rescue the growth defect caused by KKT4 RNAi (Ishii and Akiyoshi, 2020). Although the underlying molecular mechanism remains unknown, it is noteworthy that S477 is located just prior to the BRCT domain. In this study, we identified KKT8 as a putative binding partner for the KKT4 BRCT domain. It will be important to identify which kinase(s) phosphorylates the KKT8 S381 site to promote the interaction. Because kinetochore localisation of the KKT10 kinase depends on the KKT8 complex (composed of KKT8, KKT9, KKT11, and KKT12) (Ishii and Akiyoshi, 2020), it is possible that KKT10’s role in regulating the metaphase-to-anaphase transition is controlled by the KKT4^BRCT^-KKT8 interaction. These hypotheses will need to be tested in the future to better understand the mechanism of chromosome segregation in trypanosomes.

## Abbreviations

KKT4: Trypanosoma brucei KKT4
TcKKT4: Trypanosoma cruzi KKT4
LmKKT4: Leishmania mexicana KKT4

## Acknowledgments

We thank Krzysztof Kuś for his help with structural work and Richard Wheeler for providing *Leishmania mexicana* genomic DNA. We thank David Staunton for assistance with SEC-MALS experiments and Svenja Hester in the Advanced Proteomics Facility for processing mass spectrometry samples. We also thank Elspeth Garman, Midori Ishii Kanazawa, Hanako Hayashi and Lucy Cornell for comments and suggestions on the manuscript. Patryk Ludzia was supported by the Boehringer Ingelheim Fonds. Bungo Akiyoshi was supported by a Wellcome Trust Senior Research Fellowship (grant no. 210622/Z/18/Z) and the European Molecular Biology Organisation Young Investigator Program. The Department of Biochemistry NMR Facility has benefitted from funding provided by the Edward Penley Abraham Fund, the John Fell Fund and the Wellcome Trust.

## Author contributions

Patryk Ludzia purified recombinant proteins, solved crystal structures, and performed all experiments and data analysis. Edward Lowe and Gabriele Marcianò assisted in solving crystal structures of KKT4^463–645^ and *Lm*KKT4^184–284^. Shabaz Mohammed analysed crosslinking mass spectrometry data. Patryk Ludzia and Christina Redfield performed and analysed NMR experiments. Patryk Ludzia, Christina Redfield, and Bungo Akiyoshi designed experiments and wrote the manuscript.

## Materials and Methods

### Key resources table

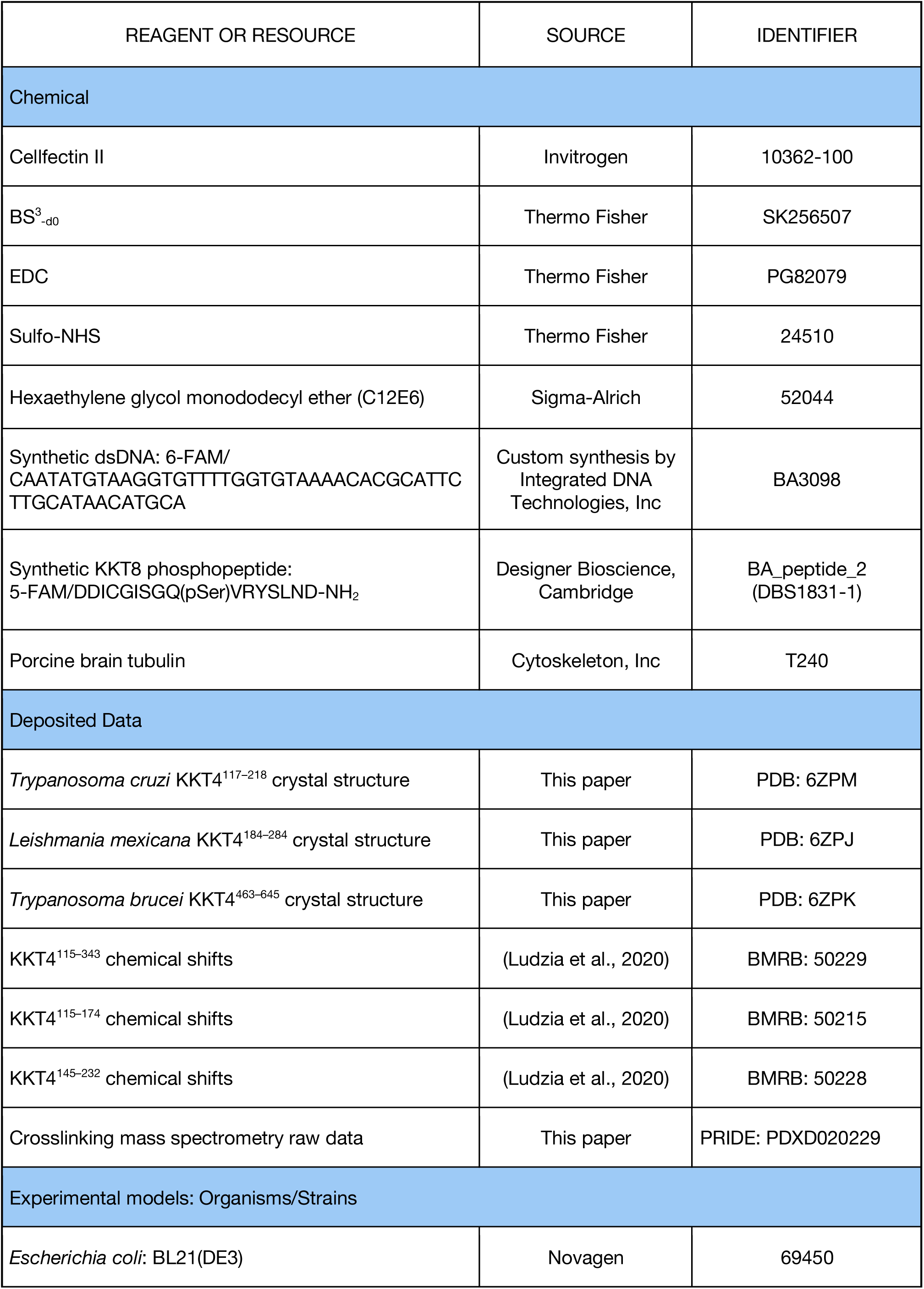
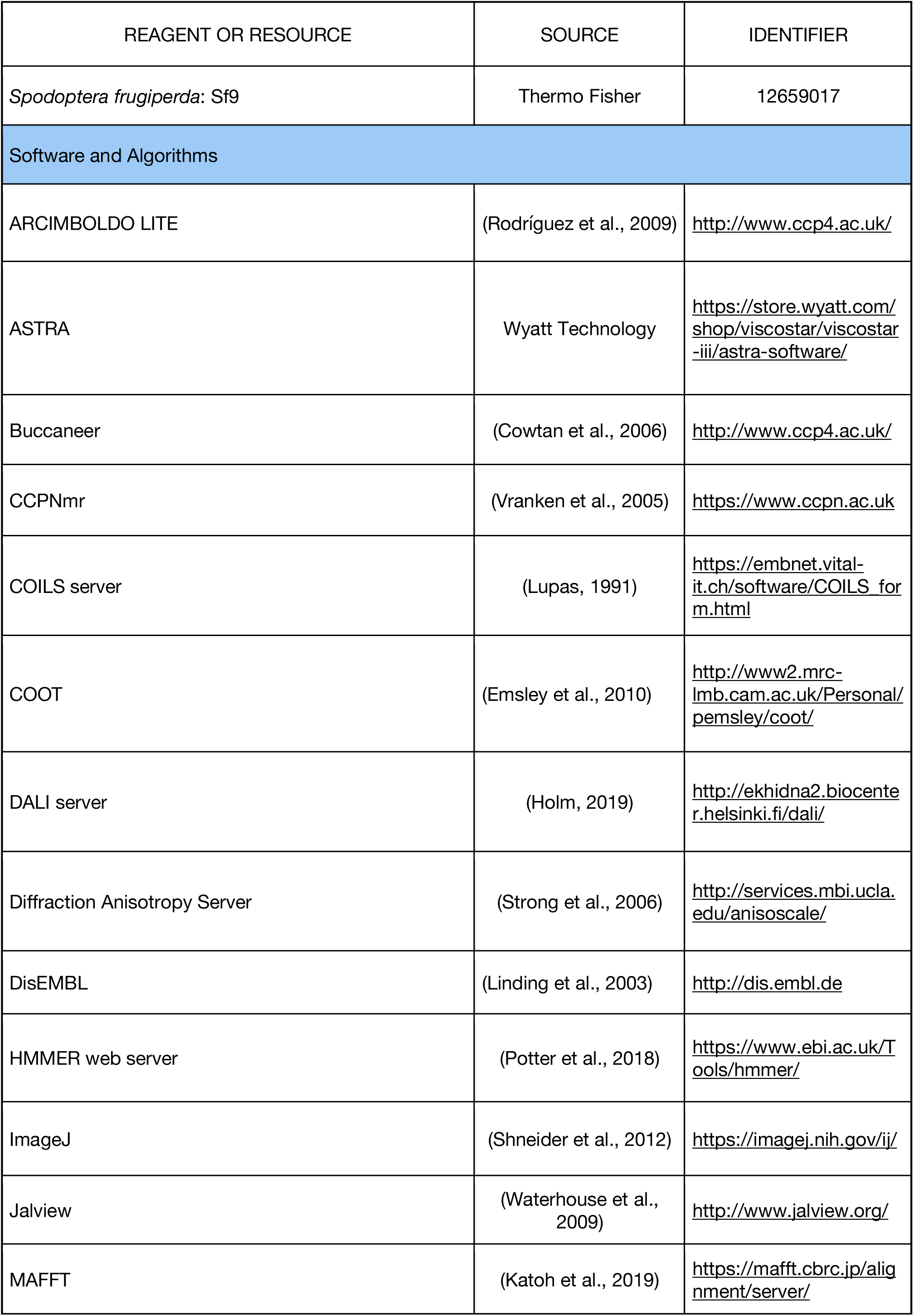
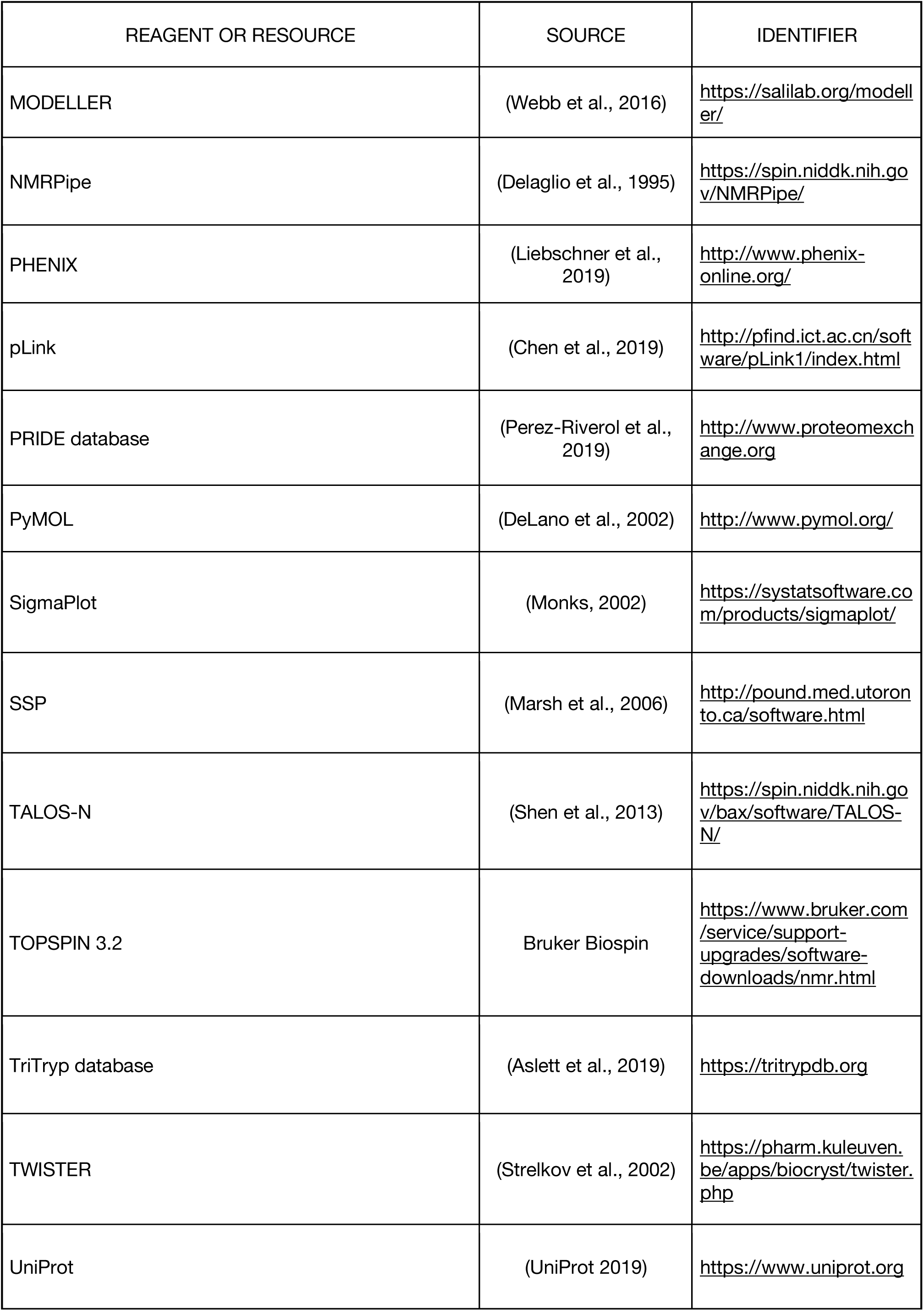
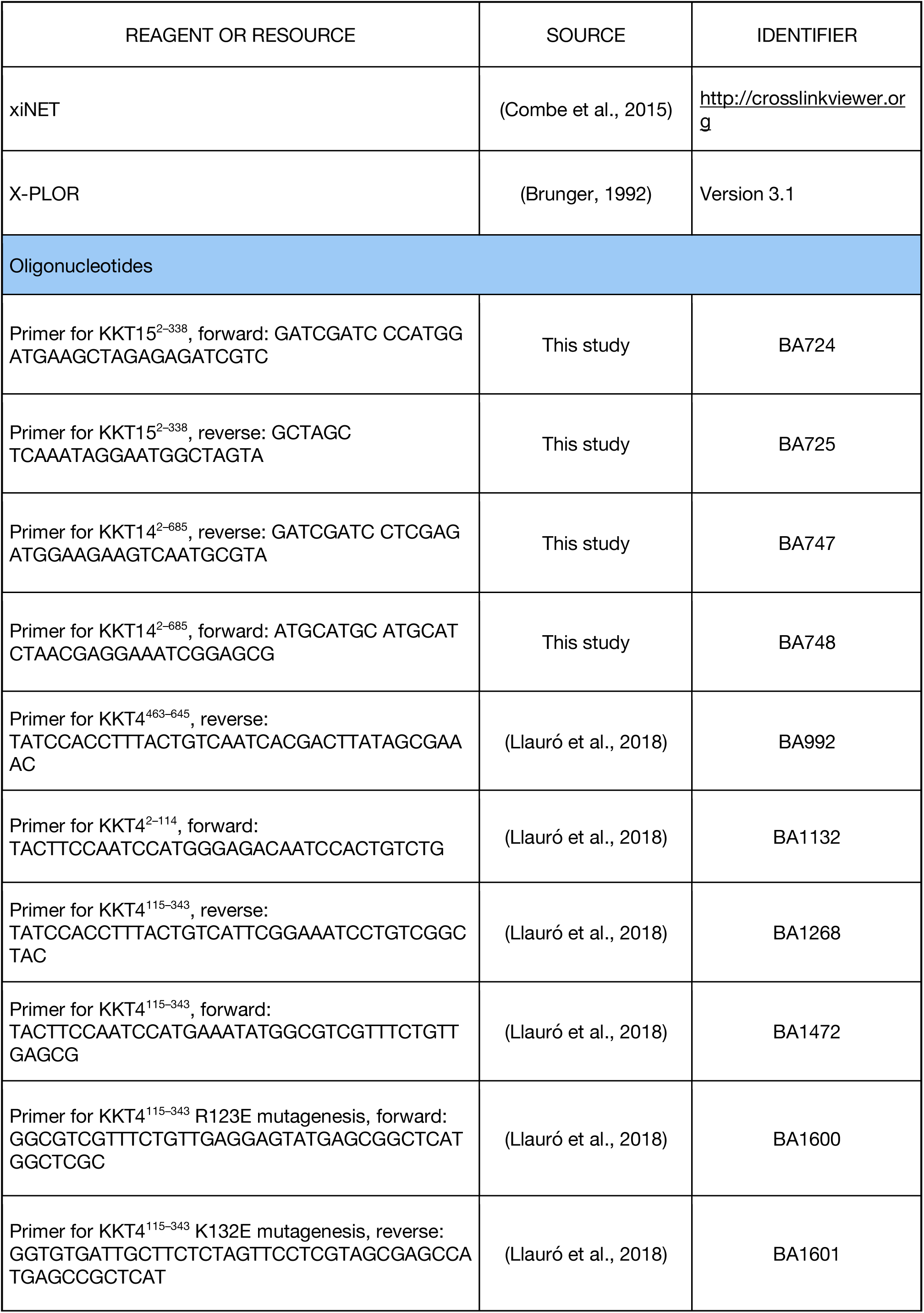
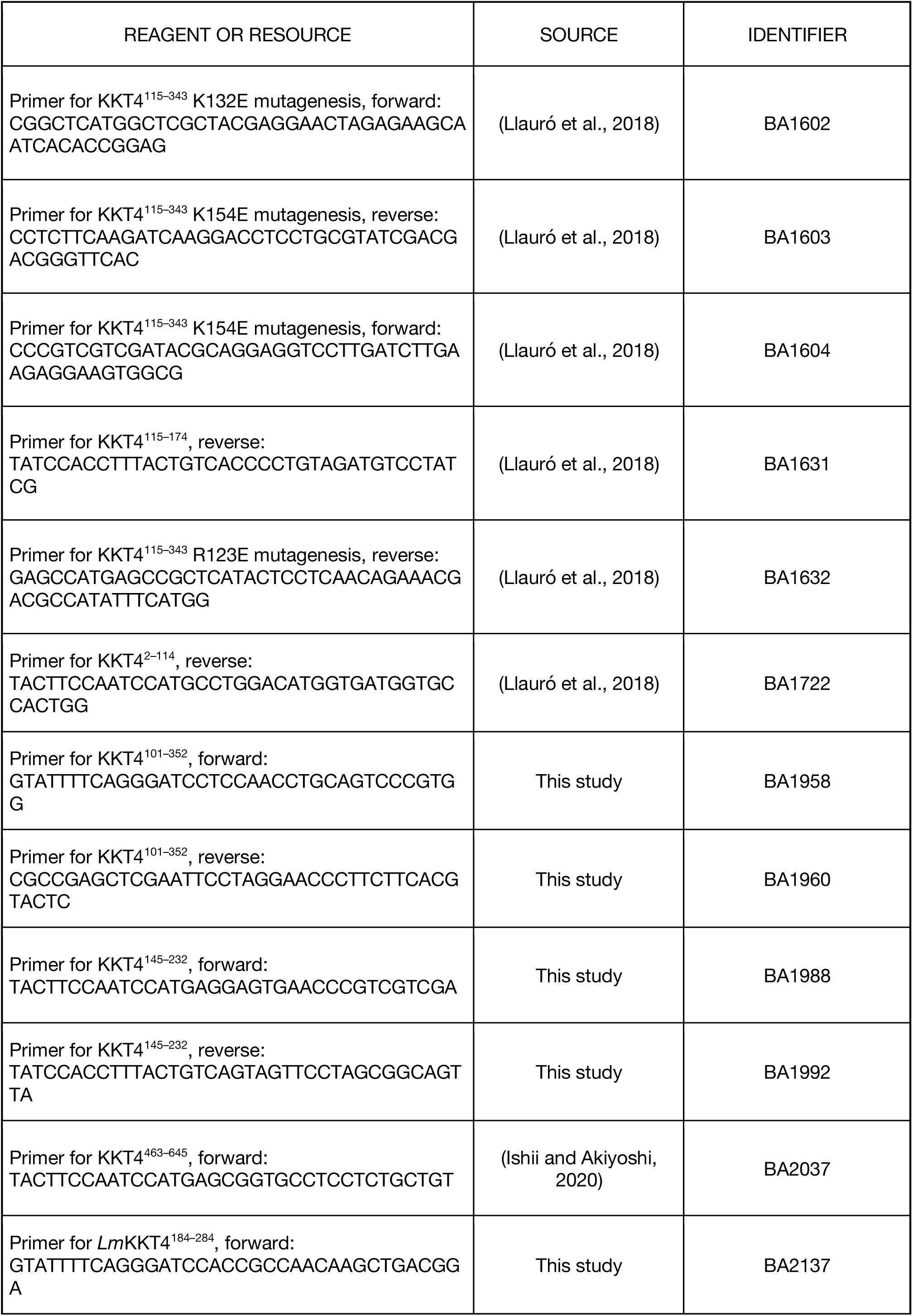
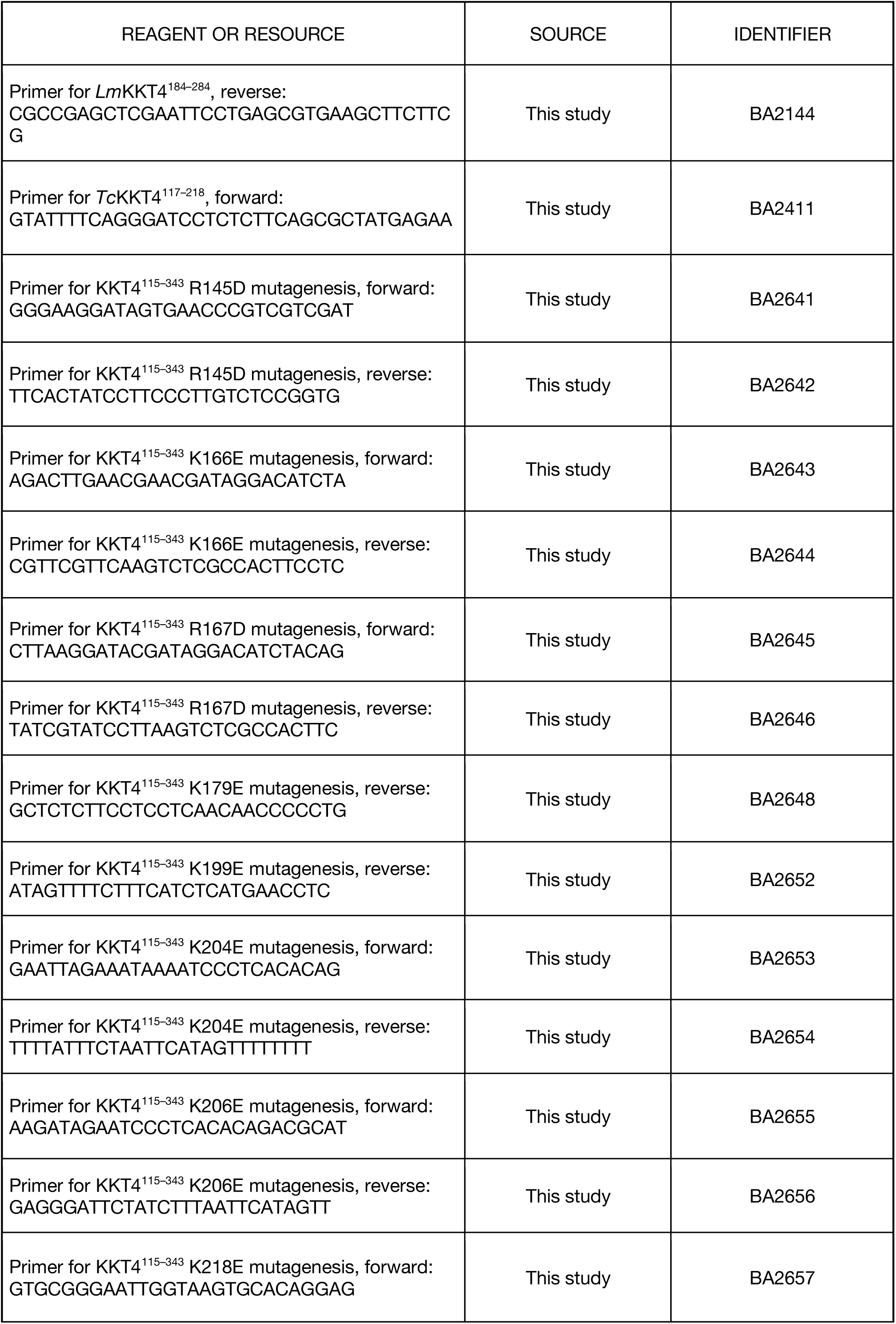
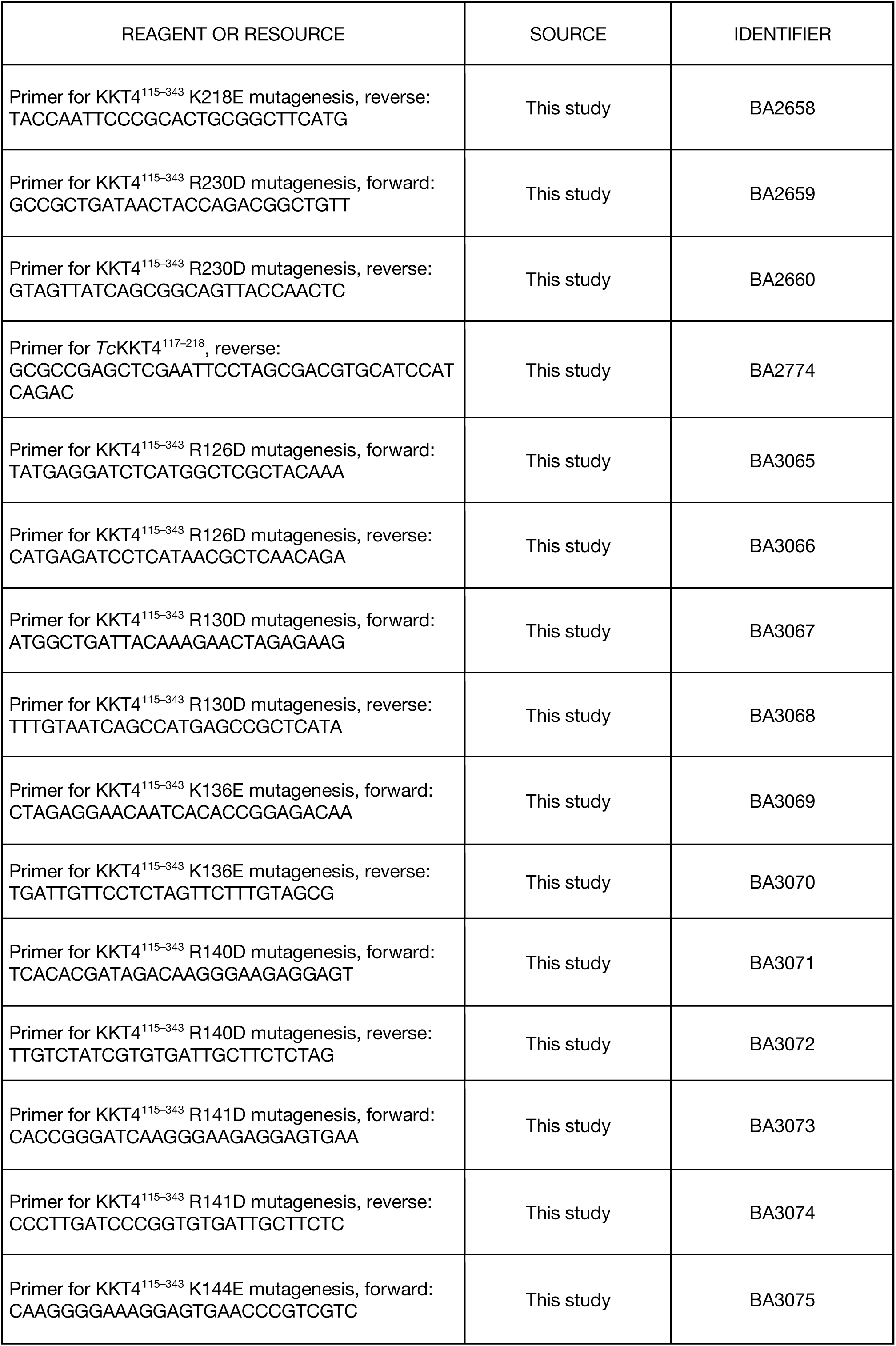
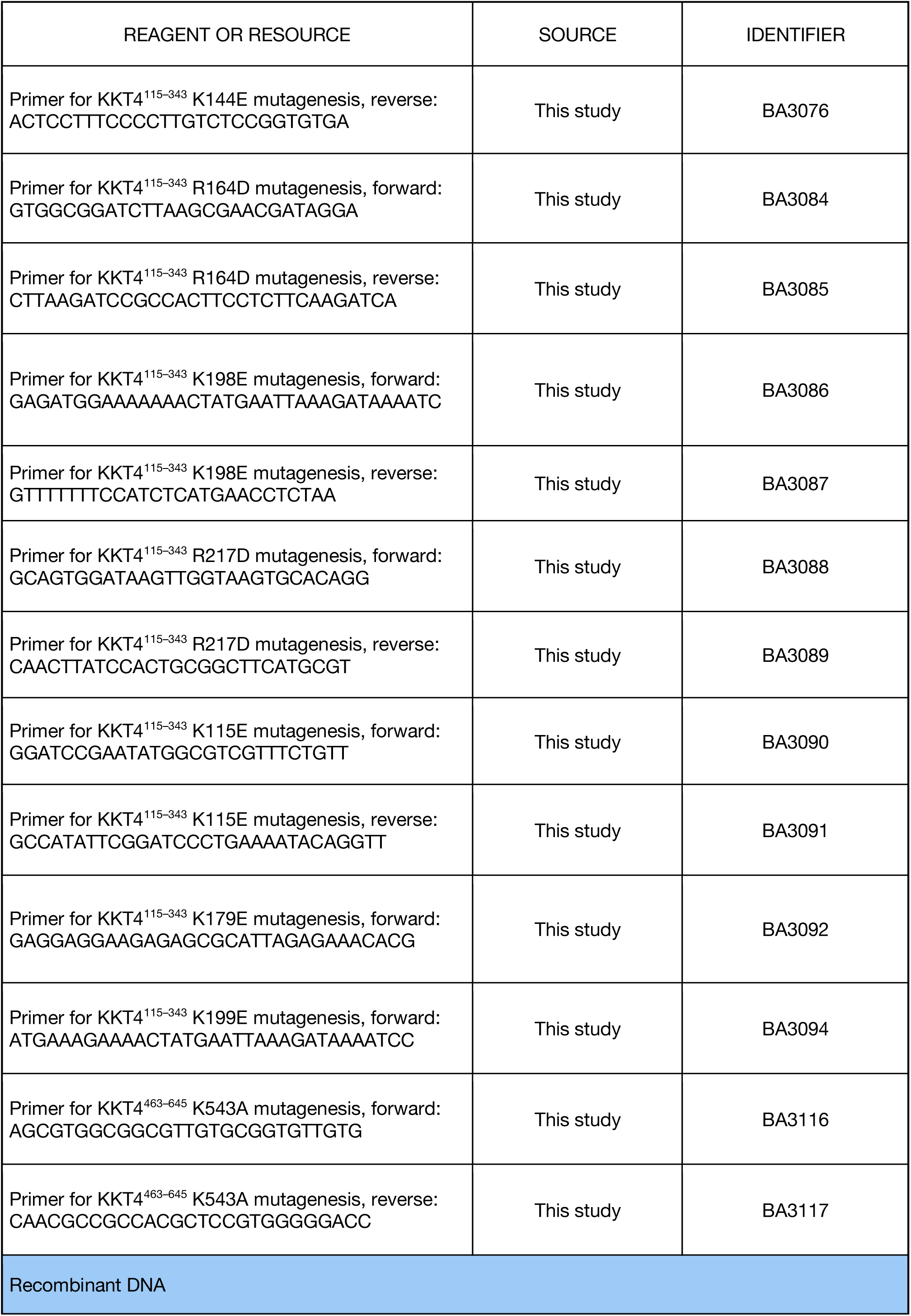
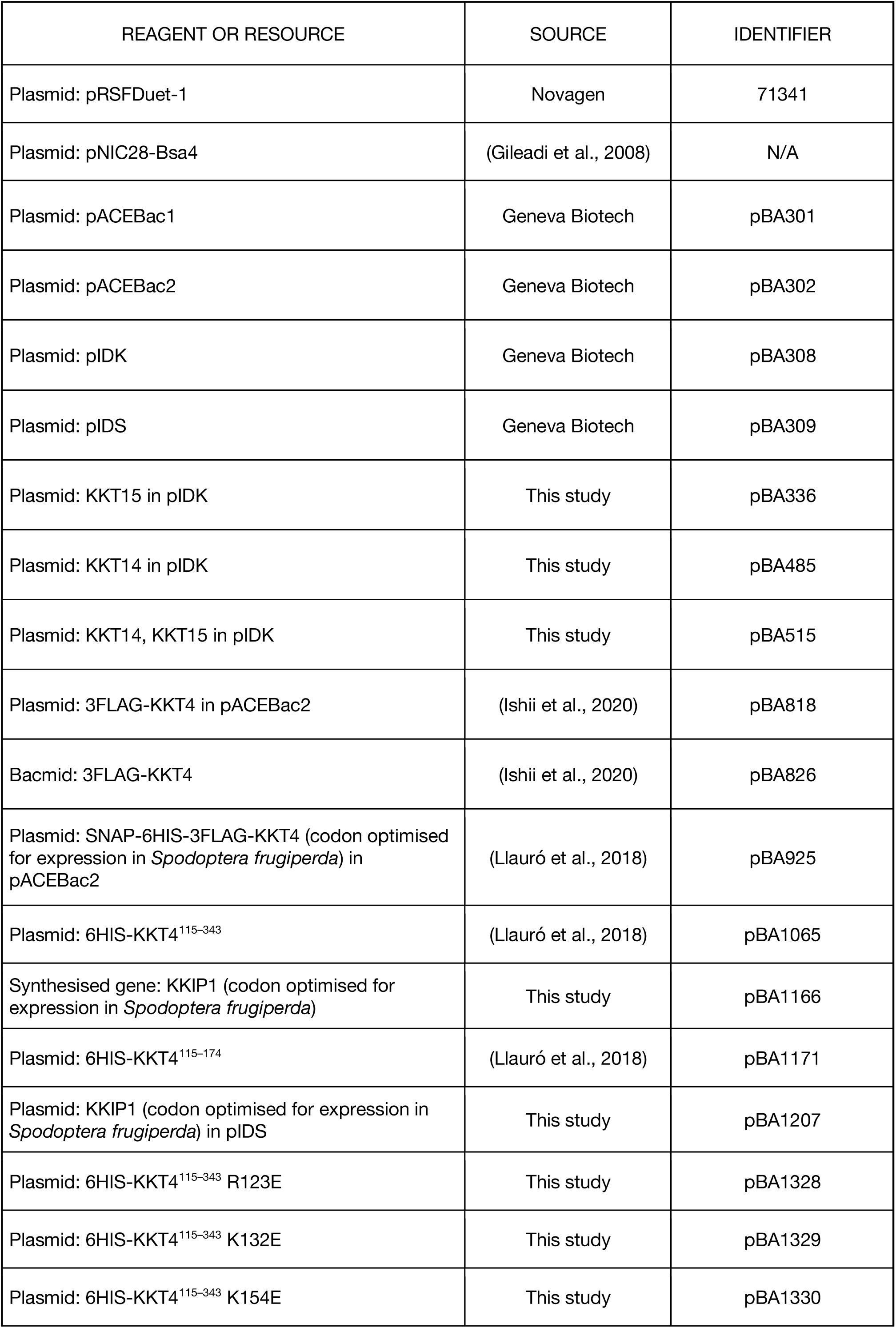
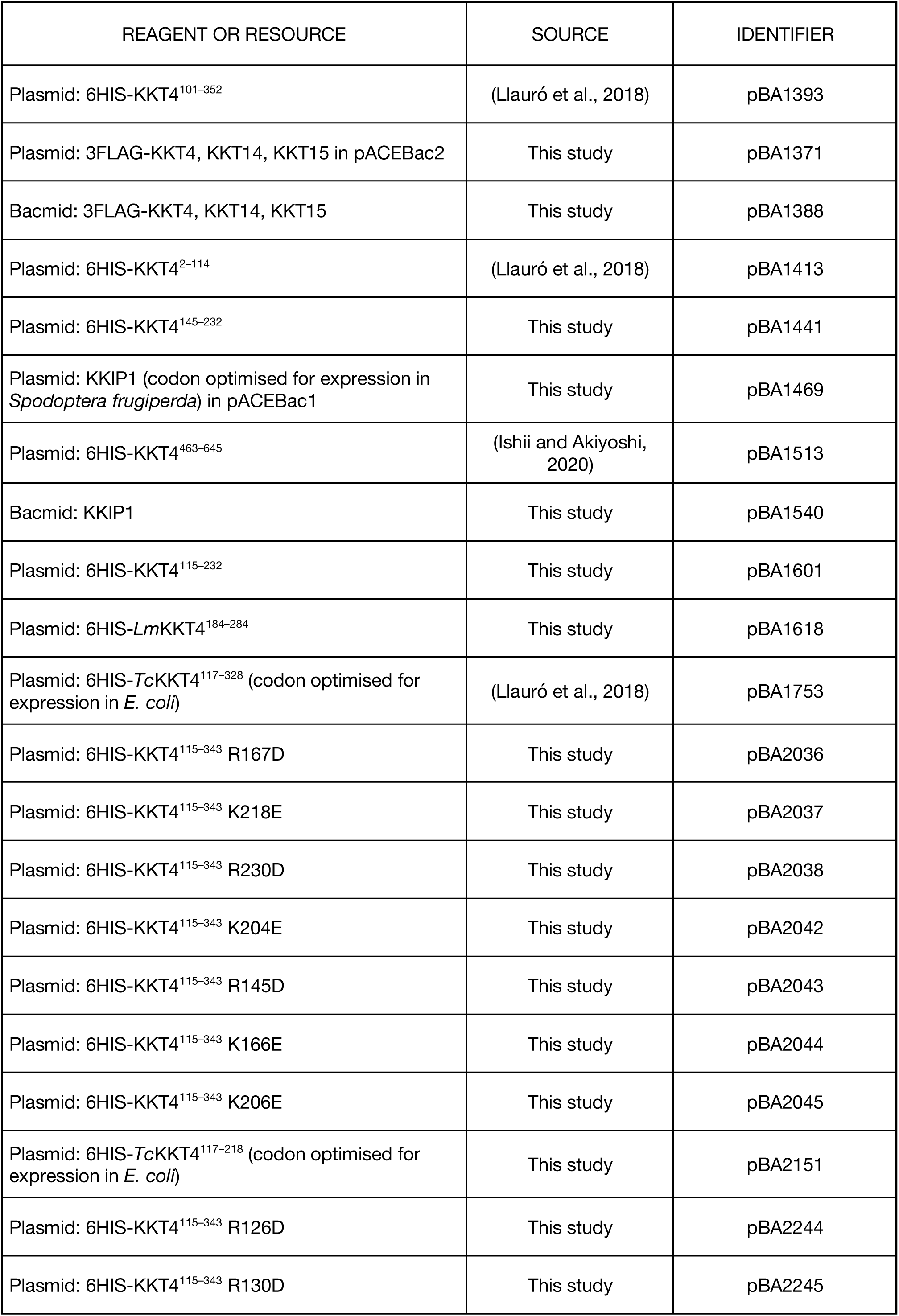
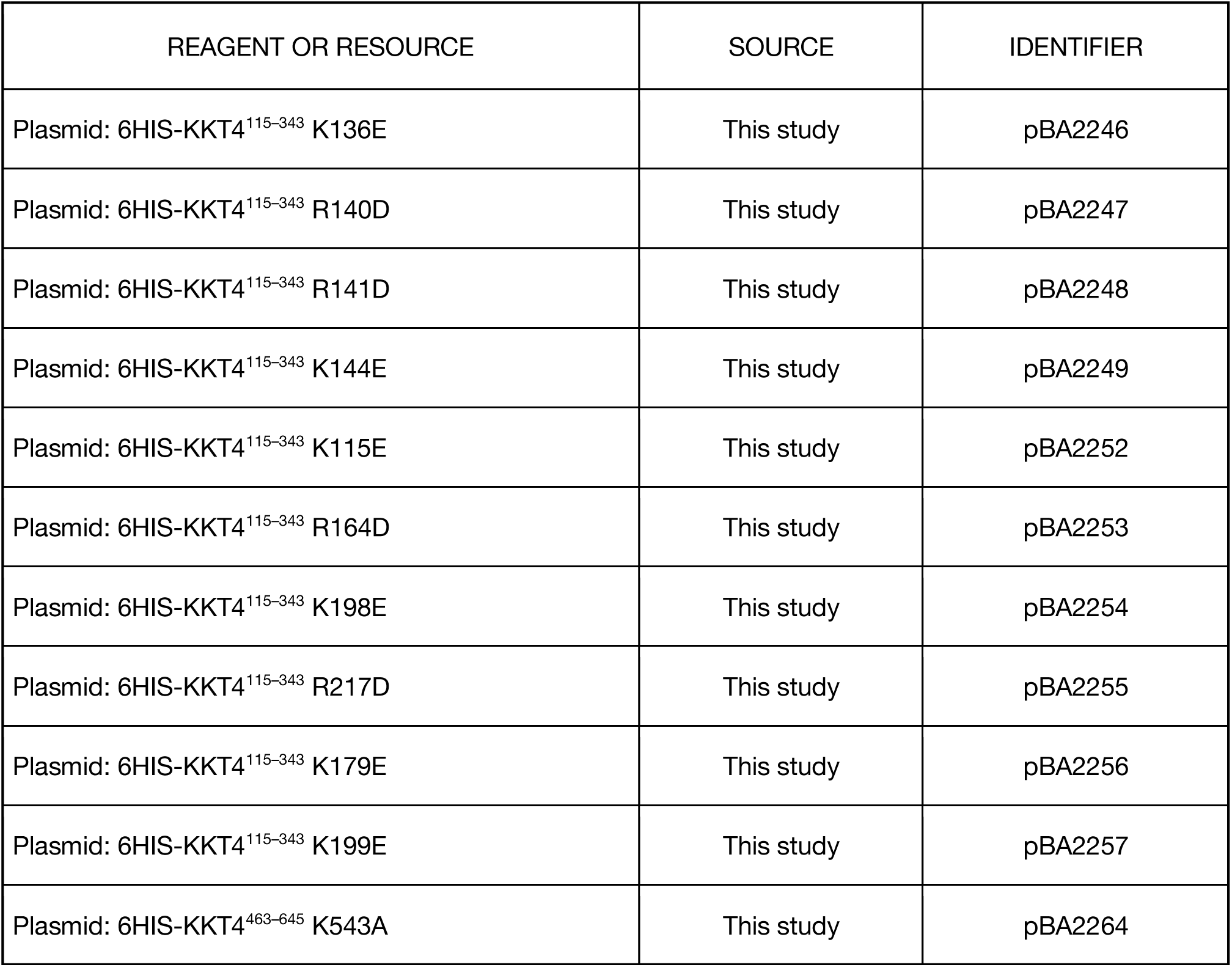

### Protein expression and purification

KKT4 fragments used in this study were amplified from *Trypanosoma brucei* genomic DNA and cloned into the pNIC28-Bsa4 expression vector using ligation-independent cloning (Gileadi et al., 2008) or cloned into RSFDuet-1 vector (Novagen) using NEBuilder HiFi DNA Assembly Kit (New England Biolabs). All constructs were sequence verified. *Lm*KKT4^184–284^ was cloned from *Leishmania mexicana* genomic DNA (kindly provided by Richard Wheeler), which contained an R218Q mutation. Due to a cloning error, which failed to place a stop codon after the *Lm*KKT4^184–284^ coding sequence, an additional 23 residues (EFELGAPAGRQACGRIMLKSNRK) from the vector were inserted at the C-terminus. *Tc*KKT4^117–218^ was cloned from synthetic *Trypanosoma cruzi* KKT4 gene fragment, codon optimised for expression in *E. coli* (Llauró et al., 2018). Point mutants of the microtubule-binding domain were created using site-directed mutagenesis using PrimeSTAR Max DNA polymerase (Takara Bio).

Transformed *E. coli* BL21(DE3) cells were inoculated into 5 ml of 2xTY medium containing 50 µg/ml kanamycin and grown overnight at 37°C. In the next morning, 1 l of 2xTY medium with 50 µg/ml of kanamycin was inoculated with 5 ml of the overnight culture and grown at 37°C with shaking (200 rpm) until the OD600 reached ∼0.7. Protein expression was induced with 0.2 mM IPTG for ∼16 hr at 16°C. Cells were spun down at 3,400 g at 4°C and resuspended in lysis buffer (50 mM sodium phosphate, pH 7.5, 500 mM NaCl, and 10% glycerol) supplemented with protease inhibitors (20 μg/ml leupeptin, 20 μg/ml pepstatin, 20 μg/ml E-64, 2 mM benzamidine, and 0.4 mM PMSF), benzonase nuclease (500 U/1 l culture), and 0.5 mM TCEP. All subsequent steps were performed at 4°C. Bacterial cultures were mechanically disrupted using a French press (1 passage at 20,000 psi) and the soluble fraction was separated by centrifugation at 48,000 g for 30 min. Supernatants were loaded on TALON beads (Takara Bio) pre-equilibrated with lysis buffer (1 ml of beads per 1 l of bacterial culture). Next, the beads were washed with lysis buffer without protease inhibitors and proteins were eluted with 50 mM sodium phosphate pH 7.5, 500 mM NaCl, 10% glycerol, 250 mM imidazole and 0.5 mM TCEP. To cleave off the His-tag, samples were incubated with TEV protease in 1:100 w/w ratio overnight while being buffer-exchanged into 50 mM sodium phosphate, 500 mM NaCl, 10% glycerol, 5 mM imidazole, and 0.5 mM TCEP by dialysis. To increase the sample purity and remove the His-tag, samples were re-loaded on TALON beads pre-equilibrated with dialysis buffer and the flow-through was collected. Next, the samples were further purified using either two-step (ion exchange and size exclusion chromatography) or one-step (size exclusion chromatography) purification. To promote binding of proteins to the ion exchange column, samples were diluted with buffer A (25 mM HEPES pH 7.5 and 0.5 mM TCEP) to achieve the final NaCl concentration of 50 mM. Ion exchange chromatography was performed using either 6 ml RESOURCE S or RESOURCE Q column (GE Healthcare) pre-equilibrated with 5% of buffer B (25 mM HEPES pH 7.5, 1 M NaCl and 0.5 mM TCEP). Proteins were eluted with a linear gradient from 5% to 100% of buffer B, concentrated using 3-or 10-kD MW Amicon concentrators (Millipore), and loaded on Superdex 75 or Superdex 200 16/60 (GE Healthcare) columns to further purify and buffer exchange into 25 mM HEPES pH 7.5, 150 mM NaCl with 0.5 mM TCEP. Fractions containing KKT4 were pooled, concentrated using a 3-or 10-kD MW Amicon concentrator (Millipore), and flash-frozen in liquid nitrogen for −80°C storage. Purification of ^15^N-labelled KKT4 proteins was done as previously described (Ludzia et al., 2020). KKT4 fragments used in microtubule co-sedimentation assays and full-length SNAP-6HIS-3FLAG-KKT4 from Sf9 insect cells were purified as described previously (Llauró et al., 2018). For crosslinking experiments with BS^3^, FLAG-KKT4 was immunoprecipitated from insect cells transfected with baculovirus prepared from pBA1388 (3FLAG-KKT4, KKT14, KKT15), whereas crosslinking with EDC/Sulfo-NHS was performed on FLAG-KKT4 that was purified from insect cells transfected with baculoviruses prepared from pBA826 (3FLAG-KKT4) and pBA1540 (KKIP1), both purified according to the protocol described in (Llauró et al., 2018). pBA1371 (3FLAG-KKT4, KKT14, KKT15 in pACEBac2) was made as follows. First, KKT15 was amplified from genomic DNA with BA724/BA725 and cloned into pIDK using NcoI/NheI, making pBA336. KKT14 was amplified from genomic DNA with BA747/BA748 into pIDK (Geneva Biotech) using XhoI/NsiI, making pBA485. Then the KKT15 expression module from pBA336 cut with PI-SceI and BstXI was ligated into pBA485 cut with PI-SceI, making pBA515. Finally, pBA818 (FLAG-KKT4 in the pACEBac2 acceptor plasmid (Ishii and Akiyoshi, 2020)) and pBA515 (KKT14, KKT15 in the pIDK donor plasmid) were fused using Cre recombinase with gentamycin-kanamycin selection, making pBA1371. pBA1469 (KKIP1 in pACEBac1) was made as follows. KKIP1 (codon optimised for expression in *Spodoptera frugiperda*) was subcloned from pBA1166 into pIDS (Geneva Biotech) using NheI/KpnI, making pBA1207. Then pBA1207 and pACEBac1 (Geneva Biotech) were fused using Cre recombinase with gentamycin-spectinomycin selection, making pBA1469. These plasmids (pBA1371 and pBA1469) were integrated into the DH10EmBacY baculoviral genome in DH10EmBacY *E. coli* cells to make bacmids (pBA1388 and pBA1540). Bacmids were purified from *E. coli* using a PureLink HiPure Plasmid Miniprep Kit (Thermo Fisher) and used to transfect Sf9 cells using Cellfectin II transfection reagent (Thermo Fisher), and baculovirus was amplified as previously described (Llauró et al., 2018).

### Size Exclusion Chromatography with Multi-Angle Light Scattering (SEC-MALS)

MALS experiments were performed during size exclusion chromatography on analytical Superdex 200 HR10/30 or Superdex 75 HR10/30 columns (GE Healthcare) equilibrated with 25 mM HEPES pH 7.5, 150 mM NaCl and 0.5 mM TCEP (for SNAP-6HIS-3FLAG-KKT4, 25 mM HEPES pH 7.5, 340 mM NaCl and 0.5 mM was used). Elution was monitored via online static light-scattering (DAWN HELEOS 8+, Wyatt Technology), differential refractive index (Optilab T-rEX, Wyatt Technology) and UV (SPD-20A, Shimadzu) detectors. Data were analysed using the ASTRA software package (Wyatt Technology).

### NMR spectroscopy

All NMR samples were prepared in 25 mM HEPES pH 7.2, 150 mM NaCl, 0.5 mM TCEP and 95% H2O/5% D2O. All NMR spectra were acquired using a 750 MHz spectrometer equipped with a Bruker Avance III HD console and a 5 mm TCI CryoProbe. All NMR data were processed using NMRPipe (Delaglio et al., 1995) and analysed using CCPN Analysis (Vranken et al., 2005).

### Analysis of chemical shifts

^1^H, ^13^C and ^15^N chemical shifts of KKT4^115–174^, KKT4^145–232^ and KKT4^115–343^ were analysed using TALOS-N (Shen and Bax, 2013) and SSP (Marsh et al., 2006) to predict secondary structure propensities.

### Backbone dynamics

The {^1^H}-^15^N heteronuclear NOE was measured for 0.2–0.5 mM samples of KKT4^115–174^, KKT4^145–232^ and KKT4^115–343^ using the TROSY-based heteronuclear NOE experiment recorded with and without ^1^H saturation for 4.5 sec at 750 MHz (Zhu et al., 2000). The data sets were acquired using 128 complex t1 increments, 96 scans per increment and with a ^15^N sweep width of 1597.444 Hz for KKT4^115–174^ and 1901.141 Hz for KKT4^145–232^ and KKT4^115–343^. 1K complex data points were recorded in the F2 dimension with a sweep width of 9259.259 Hz. Data were collected at 20°C for KKT4^115–174^ and KKT4^115–343^ and at 30°C for KKT4^145–232^. The {^1^H}-^15^N NOE was calculated as the ratio of the peak intensities in the spectra recorded with and without ^1^H saturation. Peak heights were determined using CCPN Analysis (Vranken et al., 2005). Uncertainties in the {^1^H}-^15^N NOE values were estimated from 500 Monte Carlo simulations using the baseline noise as a measure of the error in the peak heights.

### Residual Dipolar Couplings (RDCs)

Partial alignment of the KKT4^115–174^ fragment was achieved using C12E6/*n*-hexanol liquid crystals prepared as described by Rückert and Otting (Rückert and Otting, 2000). A 10% C12E6/*n-* hexanol solution was prepared in HEPES buffer and added to the protein sample to achieve the desired final concentration of 3% for KKT4^115–174^.

NMR experiments were performed at 750 MHz. ^15^N-^1^H^N^ RDCs were measured at 20°C for using BEST TROSY and BEST semi-TROSY experiments (Schulte-Herbruggen and Sorensen, 2000, Lescop et al., 2007). 128 complex points and a sweep width of 1597.444 Hz was collected in F_1_ (^15^N). Residual dipolar couplings were measured as the difference between the splitting observed in the isotropic and aligned data sets. Error bars were derived from three measurements of the RDCs.

The principle components (A_xx_, A_yy_ and A_zz_) and orientation (*ϕ*, *θ* and *ψ*) of the molecular alignment tensor were fitted to minimise the *χ*^2^ between the experimental and calculated RDCs using the *T. cruzi* X-ray coordinates. The sequence of KKT4^115–174^ was aligned with different positions in the *T. cruzi* X-ray structure in order to find the best fit of the *T. brucei* residues into the *T. cruzi* heptad repeats. Residues with {^1^H}-^15^N heteronuclear NOE values of less than 0.6 were excluded from the fitting procedure. Q values were calculated to assess the quality of the fits between experimental and calculated RDCs using the method of (Cornilescu et al., 1998).

### Crystallisation

All crystals were obtained in sitting drop vapour diffusion experiments in 96-well plates, using drops of 200 nl overall volume, mixing protein and mother liquor in a 1:1 ratio. Crystals of *Trypanosoma cruzi* (Sylvio X10) KKT4^117–218^ (10.0 mg/ml) were grown at 18℃ in Morpheus II HT-96 G3 solution (Molecular Dimensions) containing 0.1 M buffer system 4 pH 6.5, 50% v/v precipitant mix 7 and 100 mM amino acids II. Mother liquor served as a cryoprotectant. Crystals of *Leishmania mexicana* KKT4^184–284^ (13.5 mg/ml) were grown at 4℃ in solution containing 0.1 M imidazole pH 7.0 and 50% v/v MPD. Mother liquor served as a cryoprotectant. Crystals of *Trypanosoma brucei* KKT4^463–645^ (26.5 mg/ml) were grown at 4℃ in solution containing 0.1 M bis-Tris pH 5.5 and 2.0 M ammonium sulphate. Crystals were briefly transferred into mother liquor prepared with addition of 23% glycerol prior to flash-cooling by plunging into liquid nitrogen.

### Diffraction data collection and structure determination

X-ray diffraction data from *Trypanosoma cruzi* (Sylvio X10) KKT^117–218^ and *Leishmania mexicana* KKT4^184–284^ were collected at the I03 and I24 beamlines respectively, at the Diamond Light Source (Harwell, UK). The structures were solved using ARCIMBOLDO LITE optimised for coiled coils (Rodríguez et al., 2009, Rodríguez et al., 2012) followed by initial model building with BUCCANEER (Cowtan, 2006). Next, the data files were scaled to the high-resolution limit of 1.9 Å and processed using anisotropic scaling (Strong et al., 2006). Further manual model building and refinement were completed iteratively using COOT (Emsley et al., 2010) and PHENIX (Liebschner et al., 2019).

X-ray diffraction data from *Trypanosoma brucei* KKT^463–645^ were collected at the I24 beamline at the Diamond Light Source (Harwell, UK). The structure was solved using ARCIMBOLDO LITE (Rodríguez et al., 2009, Rodríguez et al., 2012) followed by initial model building with BUCCANEER (Cowtan, 2006). The further model building and refinement were completed using COOT (Emsley et al., 2010) and PHENIX (Liebschner et al., 2019).

The final refinement statistics for three structures are summarised in Table 2. All structure figures were prepared using PyMOL (DeLano, 2002). Protein coordinates have been deposited in the RCSB Protein Data Bank (http://www.rcsb.org/) with accession codes: PDB: 6ZPM (*Trypanosoma cruzi* (Sylvio X10) KKT4^117–218^), PDB: 6ZPJ (*Leishmania mexicana* KKT4^184–284^) and PDB: 6ZPK (*Trypanosoma brucei* KKT4^463–645^).

### Modelling of *T. brucei* KKT4^115–232^ and KKT4^115–645^

Homology models for the two coiled-coil regions of *T. brucei* KKT4^115–232^ were generated using Modeller 9 v24 (Webb and Sali, 2016), the X-ray structure of the *T. cruzi* KKT4^117–218^ coiled coil, and the sequence alignments of *T. brucei* and *T. cruzi* derived from the RDC data collected for KKT4^115–174^. Modeller was run using the fully automated comparative modelling mode. Random extended structures for the N- and C-termini of KKT4^115–232^ and for the inter-helix linker were generated using X-PLOR (Brünger, 1992); these represent possible conformations that these residues might sample in solution. These coordinates were merged with the two coiled-coil dimer models to generate an overall model for *T. brucei* KKT4^115–232^. The model for KKT4^115–645^ was created by merging the model for KKT4^115–232^, the X-ray structure of the BRCT domain (KKT4^473– 645^), and two random structures for disordered residues 233–472 generated using X-PLOR (Brünger, 1992). The two random structures selected to represent 233–472 where chosen from a group of ten structures to illustrate possible conformations in which the BRCT domain is in close proximity to the N-terminal coiled coil and is more distant from this region.

### Microtubule co-sedimentation assay

Taxol-stabilised microtubules were prepared by mixing 2.5 μl of 100 μM porcine tubulin (Cytoskeleton) resuspended in BRB80 (80 mM Pipes-KOH pH 6.9, 1 mM EGTA, and 1 mM MgCl_2_) with 1 mM GTP, 1.25 μl of BRB80, 0.5 μl of 40 mM MgCl_2_, 0.5 μl of 10 mM GTP, and 0.25 μl DMSO, and incubated for 20 min at 37°C. Then, 120 μl of pre-warmed BRB80 containing 12.5 μM Taxol (paclitaxel; Sigma) was added to the sample. Prior to the assay, KKT4 fragments were buffer-exchanged into BRB80 with 100 mM KCl using Zeba desalting spin columns (Thermo Fisher). For the microtubule co-sedimentation assay, 20 μl of KKT4 fragments (4 μM) were mixed with 20 μl of microtubules (2 μM) and incubated for 45 min at room temperature. For a no-microtubule control, KKT4 fragments were mixed with BRB80 with 12.5 μM Taxol. The samples were spun at 20,000 g at room temperature for 10 min, and the supernatant was collected. To the tube with a pellet, we added 40 μl of chilled BRB80 with 5 mM CaCl_2_ and incubated on ice for 5 min to depolymerise microtubules. Following the incubation, samples were boiled for 5 min before analysis by SDS-PAGE gels stained with SimplyBlue Safe Stain (Invitrogen).

### Fluorescence anisotropy assay

The DNA-binding analysis of KKT4 was performed in binding buffer (25 mM HEPES pH 7.5, 50 mM NaCl and 0.5 mM TCEP) using a 50-bp DNA probe BA3098 (∼50% GC content), which is part of the *Trypanosoma brucei* centromere CIR147 sequence (Obado et al., 2007). Prior to the assay, KKT4 proteins were buffer-exchanged into the binding buffer using Zeba spin desalting columns (Thermo Fisher). KKT4^2–645^ (0.67 µM) and KKT4^463–645^ (1 µM) samples were mixed with DNA probe in the binding buffer to a final DNA concentration of 1 nM. Next, the proteins were serially diluted in the binding buffer containing 1 nM DNA in the 2:3 v/v ratio. The binding reactions were incubated for 30 min at room temperature and fluorescence anisotropy was measured using a PHERAstar FS next-generation microplate reader (BMG LABTECH). Equilibrium dissociation constants (K_D_) were calculated by fitting the data in SigmaPlot (Monks, 2002). The phosphopeptide-binding experiments were carried out using fluorescently-labelled phosphopeptide probe. KKT4^463—645^ (170 µM) was mixed with the probe to a final peptide concentration of 100 nM and serially diluted in a 2:3 v/v ratio. Incubation of the samples and measurements were done as described above.

### Chemical crosslinking mass spectrometry (XL-MS)

Prior to the experiment, bis(sulfosuccinimidyl)suberate, BS^3^ (Thermo Fisher), crosslinker was equilibrated at room temperature for 2 hr and then resuspended to 0.87 mM in distilled water. Immediately after, 2 μl of the crosslinker was mixed with 18 μl of ∼5 μM KKT4 in 25 mM HEPES pH 8.0, 2 mM MgCl_2_, 0.1 mM EDTA, 0.5 mM EGTA-KOH, 10% glycerol, 250 mM NaCl, and 0.1% NP40. The crosslinking reaction was incubated on ice for 60 min. 1-ethyl-3-(3-dimethylaminopropyl)carbodiimide hydrochloride (EDC) and *N*-Hydroxysulfosuccinimide sodium salt (Sulfo-NHS) were resuspended in distilled water to 4 mM and 10 mM respectively. Immediately after, 1 μl of EDC and 1 μl of Sulfo-NHS were mixed with 18 μl of ∼5 μM KKT4. The crosslinking reaction was incubated at room temperature for 60 min. Following the incubation, all the crosslinking samples were boiled for 10 min and resolved on a NuPAGE 4–12% gradient polyacrylamide gel (Invitrogen). Gels were stained using SimplyBlue (Invitrogen) and bands corresponding to crosslinked KKT4 were cut out and subjected to mass spectrometry. Gel bands were destained by cycles of incubation with 50% acetonitrile in 100 mM triethylammonium bicarbonate (TEAB) and 100% acetonitrile. Subsequently, dried gel bands were incubated with 10 mM TCEP in 100 mM TEAB for 30 min at room temperature followed by centrifugation. Supernatant was removed. Subsequently, 50 mM 2-chloroacetamide in 100 mM TEAB was added to the gel and incubated for 30 min at room temperature in the dark. Gel bands were subsequently washed with 100% acetonitrile 3 times followed by addition of 100 ng of trypsin and digested overnight at 37°C. Subsequently, the supernatant was extracted. The gel pieces were then washed with an 5% formic acid and then acetonitrile. Both washes were combined with the initial supernatant and the mixture was then dried down.

Peptides were resuspended in 5% formic acid and 5% DMSO and analysed by LC-MS with an Ultimate 3000 UHPLC system (Thermo Fischer Scientific) coupled to an QExactive mass spectrometer (Thermo Fischer Scientific) through an EASY-Spray nano-electrospray ion source (Thermo Fischer Scientific). The peptides were trapped on a C18 PepMap 100 pre-column (300 μm i.d. x 5 mm, 100Å, Thermo Fisher Scientific) using solvent A (0.1% formic acid in water). The peptides were separated on an in-house packed analytical column (75 μm i.d. x 50 cm packed with ReproSil-Pur 120 C18-AQ, 1.9 μm, 120 Å, Dr. Maisch GmbH) using a linear gradient (length: 45 min, 15% to 50% solvent B (acetonitrile with 0.1% formic acid)). Acquisition was performed in data-dependent mode (DDA). Full scan MS spectra were acquired in the Orbitrap (scan range 350–1,500 m/z, resolution 70,000, AGC target 3 x 10^6^, maximum injection time 50 ms) followed by 10 MS/MS events at 30% NCE (resolution 17,500, AGC target 5 x 10^4^, maximum injection time 120 ms, isolation window 1.5 m/z) with first fixed mass at 180 m/z. Charge exclusion was selected for unassigned 1+ and 2+ ions.

MS data were converted into mgf format using pParse and searched by the pLink software (Chen et al., 2019) (version 1 for BS^3^ dataset and version 2 for EDC dataset) using FASTA databases containing KKT1–4, 6, 7–11, 14, 15, 20, and α/β tubulins without KKIP1 (for BS^3^ dataset) or with KKIP1 (for EDC dataset). Search parameters were as follows: maximum number of missed cleavages = 2, fixed modification = carbamidomethyl-Cys, variable modification 1 = Oxidation-Met, variable modification 2 = Glu to pyro-Glu. Crosslinks that have score < 1 x 10^-7^ were visualised using xiNET (Combe et al., 2015) (Supplemental Table S4 and S5).

### Interaction of the microtubule-binding and BRCT domains

^1^H-^15^N BEST TROSY spectra were collected at 20°C for 0.2 mM KKT4^115–174^, KKT4^115–232^, KKT4^145–232^ and KKT4^115–343^ alone and in the presence of 0.22 mM KKT4^463–645^. Following addition of KKT4^463–645^ the samples were incubated at room temperature for ∼30 min. Interaction was monitored by comparing peak positions; the combined chemical shift changes of ^1^H^N^ and ^15^N are reported in Hz in Supplemental Figure 9.

### Multiple sequence alignment

Protein sequences and accession numbers for KKT4 and KKT8 homologues used this study were retrieved from the TriTryp database (Aslett et al., 2010), UniProt (UniProt, 2019), or a published study (Butenko et al., 2020). Searches for homologous proteins were done using BLAST in the TriTryp database (Aslett et al., 2010) or Jackhmmer on the UniProtKB proteome database using a default setting (HMMER web version 2.24) (Potter et al., 2018). Multiple sequence alignment was performed with MAFFT (L-INS-i method, version 7) (Katoh et al., 2019) and visualised with the Clustalx coloring scheme in Jalview (version 2.10) (Waterhouse et al., 2009).

## Data availability

All data generated during this study are included in the manuscript and supplementary data. Protein coordinates have been deposited in the RCSB Protein Data Bank (http://www.rcsb.org/) with accession codes PDB: 6ZPK (*Trypanosoma brucei* KKT4^463–645^), PDB: 6ZPM (*Trypanosoma cruzi* Sylvio X10 KKT4^117–218^) and PDB: 6ZPJ (*Leishmania mexicana* KKT4^184–284^). The chemical shift assignments for KKT4 have been deposited in the BioMagResBank (http://www.bmrb.wisc.edu) under the accession numbers 50215, 50228 and 50229. All raw files relating to crosslinking mass-spectrometry have been deposited to the ProteomeXchange Consortium via the PRIDE partner repository (Perez-Riverol et al., 2019) with the dataset identifier PXD020229 (Username, Password: 41cy6ucV).

Further information and requests for reagents may be directed to, and will be fulfilled by, the Lead Contact Bungo Akiyoshi.

## Supplemental Information

### Supplemental Tables

Supplemental Table 1: List of DALI search hits for the *Trypanosoma cruzi* KKT4^117–218^ structure

Supplemental Table 2: List of DALI search hits for the *Leishmania mexicana* KKT4^184–284^ structure

Supplemental Table 3: List of DALI search hits for the *Trypanosoma brucei* KKT4 BRCT structure

Supplemental Table 4: List of BS^3^-mediated crosslinks of KKT4 detected by mass spectrometry

Supplemental Table 5: List of EDC-mediated crosslinks of KKT4 detected by mass spectrometry

## Supplemental Figure legends

**Supplemental Figure 1.**
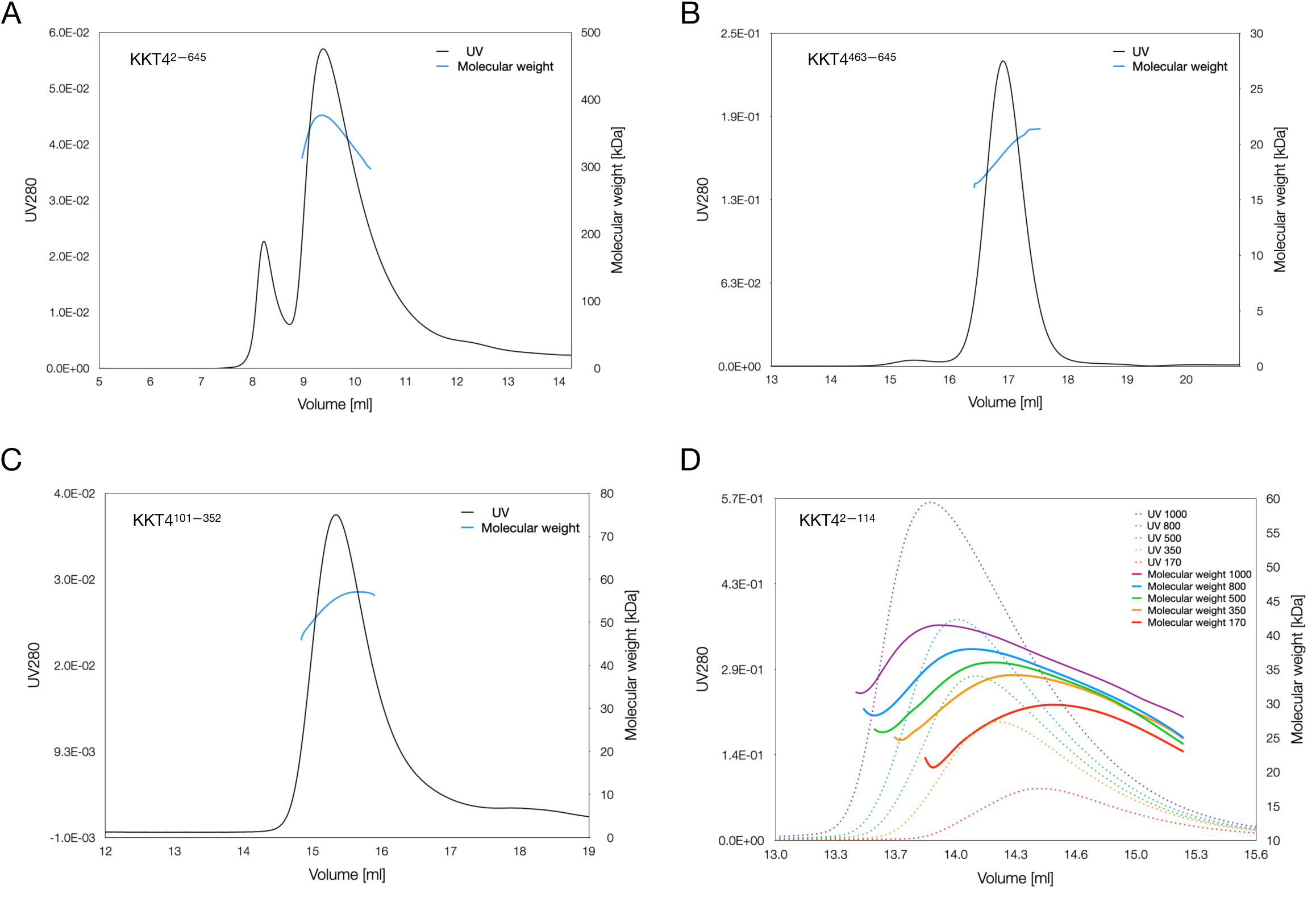
SEC-MALS analysis of KKT4 fragments. (A) SEC-MALS elution profile of SNAP-3FLAG-6HIS-KKT4^2–645^. The UV signal is plotted against the elution volume. Molecular weight is indicated as a blue line in the middle of the main peak. The molecular weight is estimated to be around 350 kDa, indicating a tetramer. (B) SEC-MALS elution profile of KKT4^463–645^. The molecular weight is estimated to be around 19 kDa, indicating a monomer. (C) SEC-MALS elution profile of KKT4^101–352^. The molecular weight is estimated to be around 55 kDa, indicating a dimer. (D) SEC-MALS elution profile of KKT4^2–114^ samples at different concentrations. The lower the protein concentration, the larger the elution volume of the protein. This suggests that oligomerisation states of KKT4^2–114^ are concentration dependent, ranging from tetramer (high concentration) to trimer (low concentration). Superdex 200 HR10/30 was used for KKT4^2–645^, KKT4^463–645^, KKT4^101– 352^ and Superdex 75 HR10/30 was used for KKT4^2–114^.

**Supplemental Figure 2.**
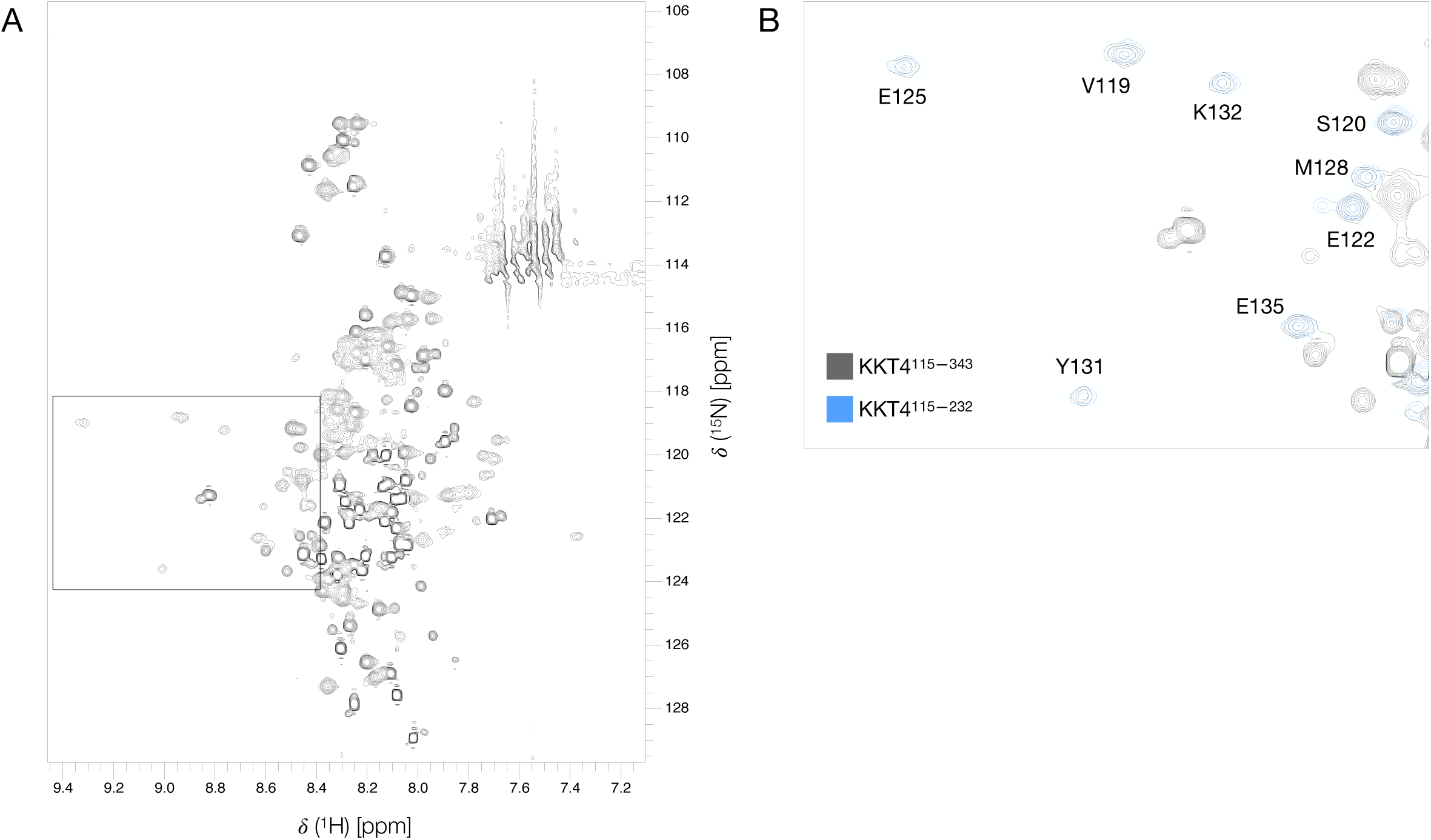
1H-^15^N BEST-TROSY spectra of KKT4^115–343^. (A) 750 MHz ^1^H-^15^N BEST-TROSY spectrum of KKT4^115–343^. The spectrum is contoured so that both weak and strong peaks are visible. Peaks in the region of 111–114 ppm and upfield of ∼7.6 ppm are artefacts in the BEST-TROSY arising from incomplete cancellation of signals from the side chain amides of asparagine and glutamine. The area outlined with a dashed box is expanded in (B). (B) Overlay of a small region of the ^1^H-^15^N BEST-TROSY spectra of KKT4^115–343^ and KKT4^115–232^. N-terminal peaks overlay well between the two constructs, suggesting that the structure of the KKT4 N-terminus is similar in both fragments.

**Supplemental Figure 3.**
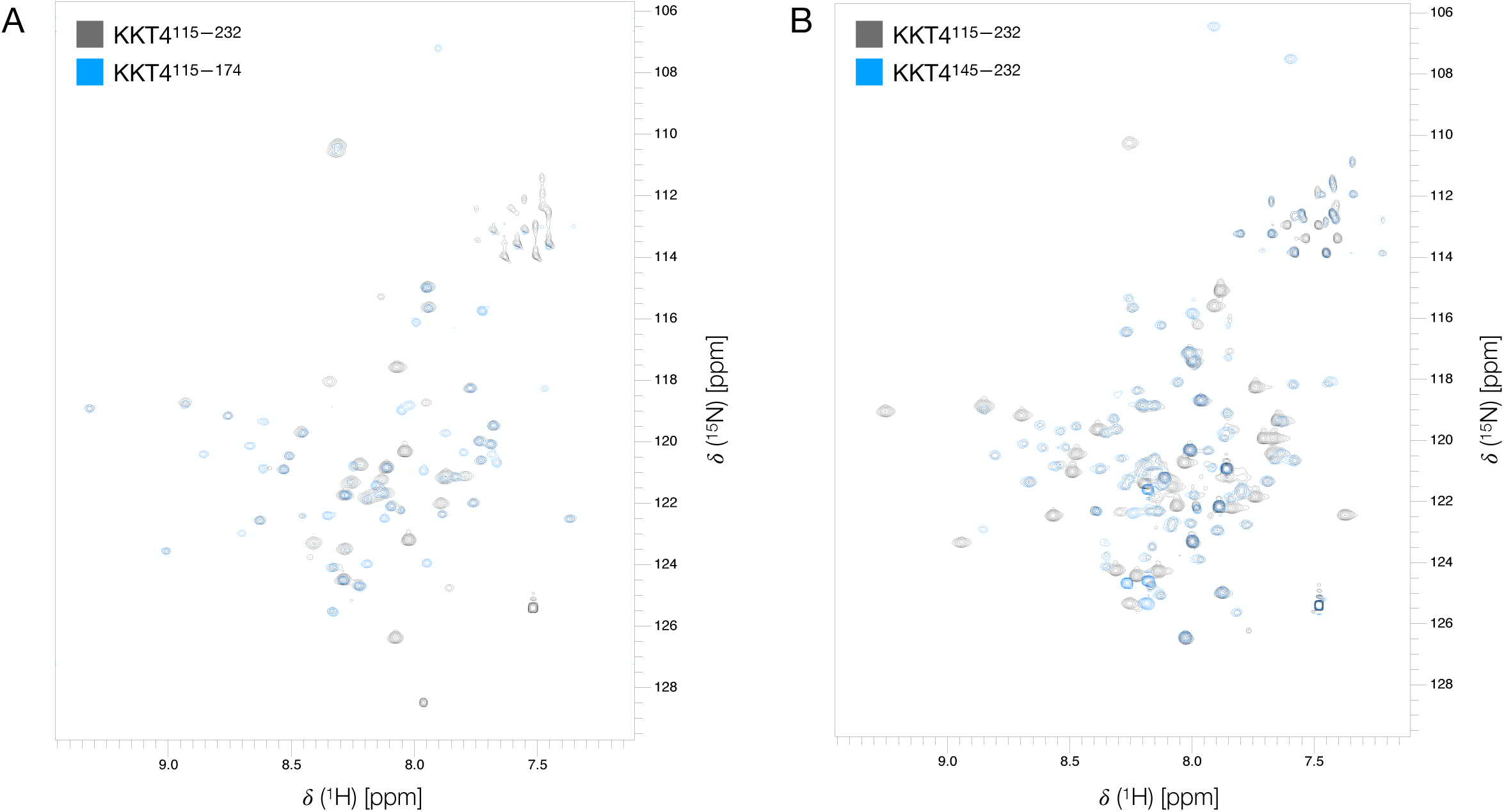
1H-^15^N BEST-TROSY overlays of KKT4 fragments show structural similarity between different KKT4 constructs. Overlay of 750 MHz ^1^H-^15^N BEST-TROSY spectra of KKT4^115–232^ and KKT4^115–174^ (A) or KKT4^115–232^ and KKT4^145–232^ (B). Most of the peaks in the spectra of KKT4^115–174^ and KKT4^145–232^ overlay with peaks observed for KKT4^115–232^, indicating that the dissection approach used was justified. Peaks that do not overlay arise from the different C-terminal sequence in KKT4^115–174^ and N-terminal sequence in KKT4^145–232^. Some peaks visible in the spectra of the shorter KKT4 fragments are not visible in the longer construct, KKT4^115–232^.

**Supplemental Figure 4.**
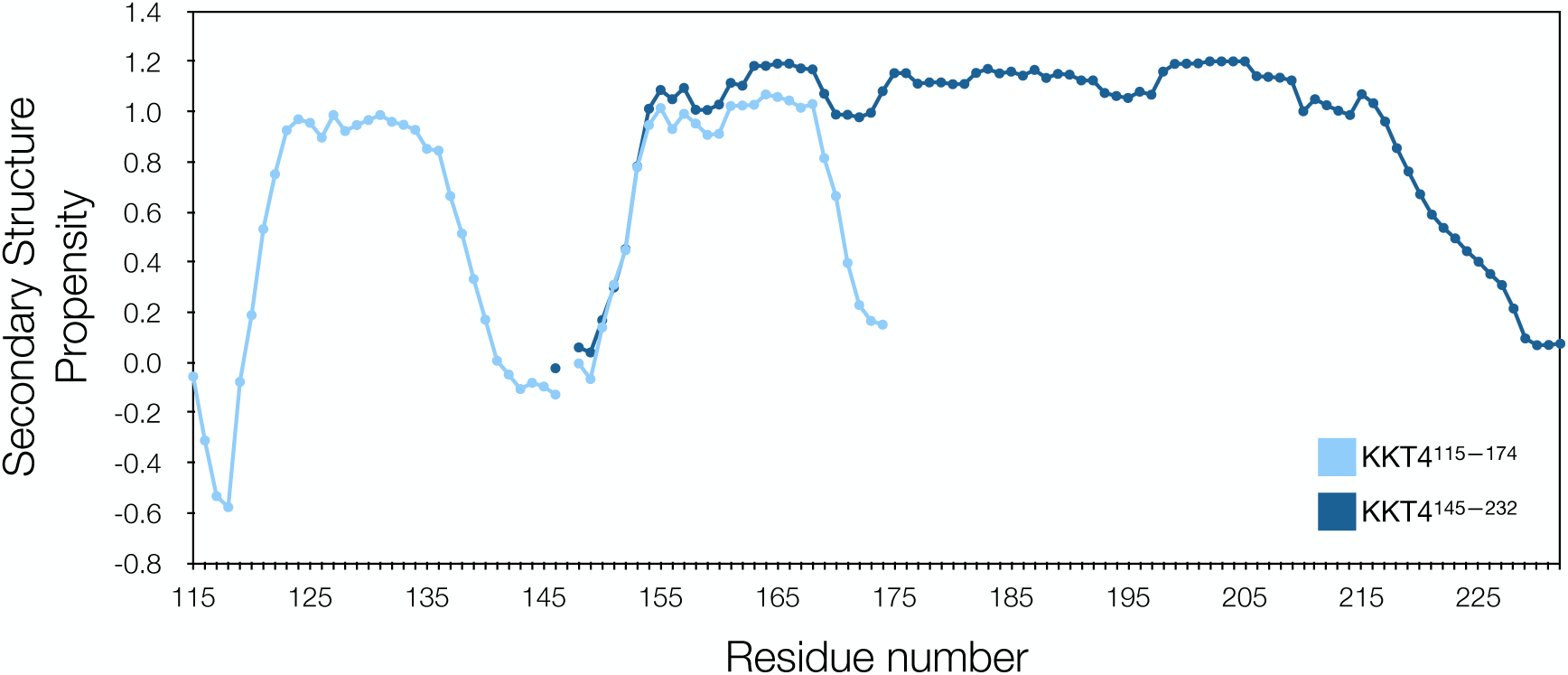
SSP analysis of KKT4^145-232^ and KKT4^115-174^. Secondary structure propensity (SSP) scores (Marsh et al., 2006) for KKT4^145–232^ and KKT4^115–174^ plotted against the protein sequence show helical secondary structure propensity in both constructs. Residues 153–223 in KKT4^145–232^ and 121–138 and 153-171 in KKT4^115–174^ have helical propensity greater than 0.5. H139 has a helical propensity of 0.33, while D151 and T152 have helical propensities of 0.31 and 0.45. H171 and L172 have helical propensities of 0.40 and 0.23. Residues 140 to 150 have propensity for either helix or sheet of less than 0.2. The gaps in the graph correspond to residues for which the NMR peaks could not be assigned.

**Supplemental Figure 5.**
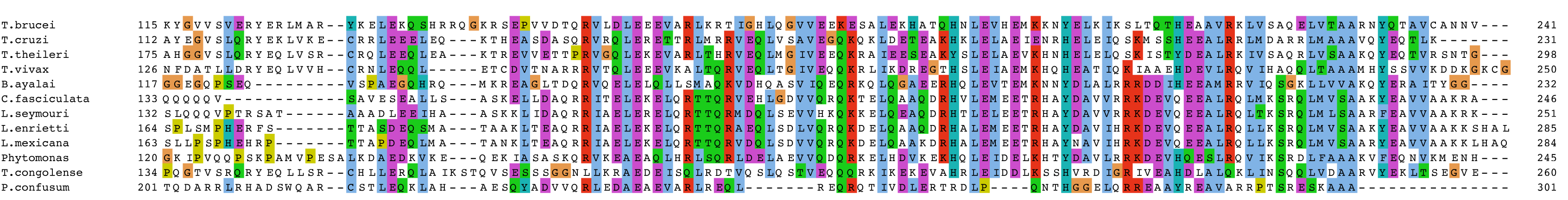
Multiple sequence alignment of N-terminal part of KKT4 microtubule-binding domain. KKT4 protein sequences from several kinetoplastid species were aligned using MAFFT (Katoh et al., 2019) and visualised with the CLUSTALX colouring scheme in Jalview (Waterhouse et al., 2009).

**Supplemental Figure 6.**
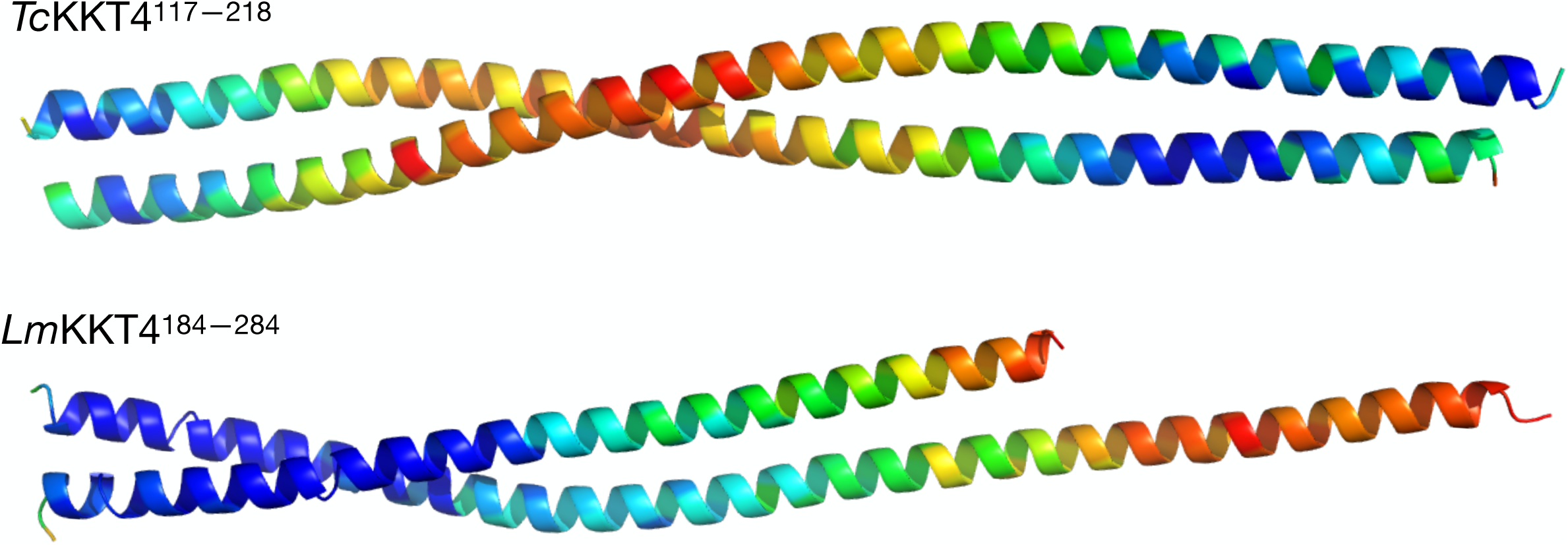
Ribbon model of *Tc*KKT4^117–218^ and *Lm*KKT4^184–284^ crystal structures colour-coded by B-factors. B-factors for Cα atoms have been represented using a blue to red spectrum indicating low to high values, respectively. The figure was rendered using PyMol (DeLano, 2002).

**Supplemental Figure 7.**
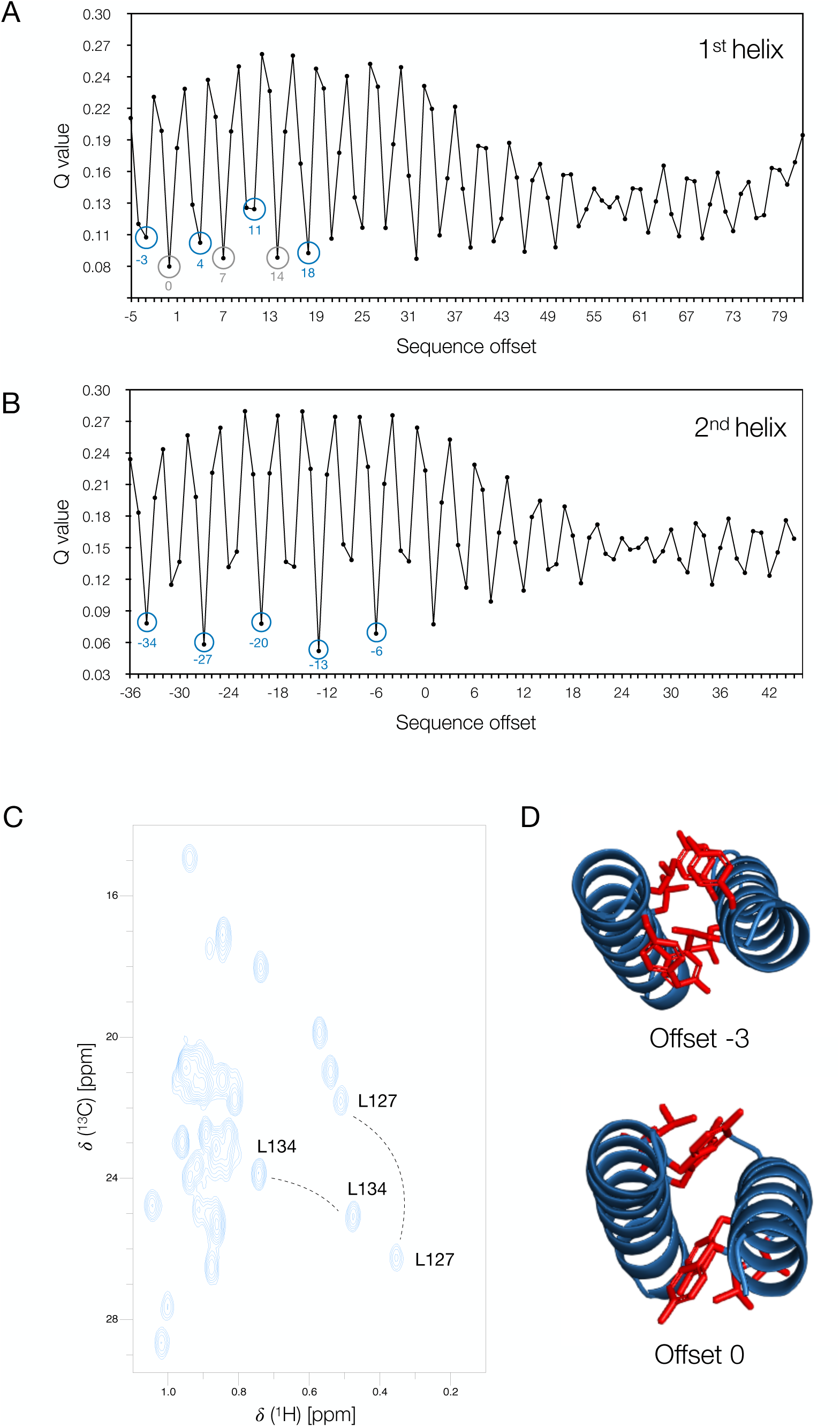
Fitting the experimental RDC data using the *Tc*KKT4^117–218^ crystal structure. The experimental RDCs for residues 126–135 from the 1^st^ helix (A) and 154–170 from the 2^nd^ helix (B) from *T. brucei* KKT4^115–174^ were fitted to the calculated RDCs based on the *T. cruzi* KKT4 X-ray structure. The relative alignment of the *T. brucei* and *T. cruzi* sequences was varied and the quality of the fit (Q value) of the experimental to predicted RDCs was measured. The Q value obtained is plotted as a function of the sequence offset (the number that is added to the *T. brucei* sequence before comparison with RDCs calculated for the *T. cruzi* sequence). For an ideal coiled-coil structure, the Q value should vary over a heptad repeat but the fits should be identical from one heptad repeat to the next. In contrast, for a more variable structure, where the interaction of the two helices varies along the length of the coiled coil, the quality of the fit may change along the sequence. For both helices, several good fits were found when the residues from *T. brucei* were aligned with the first half of the *T. cruzi* sequence, corresponding to the regular coiled-coil structure found between residues 121 and 176. Poorer agreement was obtained when the *T. brucei* helices were aligned with the C-terminal half of the *T. cruzi* structure, which shows less supercoiling. For the 2^nd^ helix (residues 154–170, shown in (B)) low Q values are found, at intervals of 7 residues (indicated by blue circles), when the residues from *T. brucei* were aligned with the residues in the N-terminal half of the *T. cruzi* sequence. The sequence alignment of these two proteins (Figure 3A) in the region of the 2^nd^ helix suggests an offset of −6 residues, which is consistent with the fits of the RDC data, but offsets of −13, −20, −27 and −34 also produce good fits (indicated by blue circles). Predictions using the COILS server suggest that the hydrophobic residues identified as occupying the *a*/*d* positions in the *T. brucei* heptad align with the *a/d* residues in the *T. cruzi* structure for offsets of −6, −13, etc. For the 1^st^ helix (residues 126–135, shown in (A)) low Q values are found for two sets of residues, at intervals of 7 residues. The lowest Q values are for sequence offsets of 0, 7, 14 (indicated by grey circles) but low Q values are also found for offsets of −3, 4, 11, 18 (indicated by blue circles); this type of degeneracy has been observed previously for the coiled-coil domain of cGK1α^9-44^ (Schnell et al., 2005). (C) The ^1^H-^13^C HSQC of KKT4^115-174^ shows that both of the methyl groups of L127 and one of the methyl groups from L134 have ^1^H peaks that are shifted upfield (to the right) of the peaks arising from most other L/V/I residues (around 0.7–1.05 ppm). These upfield shifts are due to the interaction of L127 and L134 with the aromatic rings of Y124 and Y131 and can be used to resolve the degeneracy between the two possible sequence offsets. Homology models for the 1^st^ helix of *T. brucei* KKT4^115– 174^ have been built with Modeller using the *T. cruzi* X-ray structure and sequence alignments based on sequence offsets of −3 and 0 residues; these are shown in (D) with the side chains of Y124/L127/Y131/L134 shown as red sticks. The −3 sequence offset places Y124/Y131 and L127/L134 in positions *a* and *d* of the heptad, respectively, where they interact closely. The 0 sequence offset places these pairs of residues in positions *d* and *g*; L127/L134 in the *g* position are not closely packed at the inter-helix interface. ^1^H chemical shifts for the L127 and L134 methyl groups were then predicted for the two alternative coiled-coil models. Only the homology model generated with a sequence offset of −3 residues predicts upfield shifts for the methyl groups of L127 and L134 due to their close proximity to Y124 and Y131 within the same helix and between helices in the packing interface. The sequence alignment of the *T. brucei* and *T. cruzi* proteins (Figure 3A) in the region of the 1^st^ helix suggests an offset of −3 residues, which is consistent with the RDC data and chemical shift analysis of the homology models.

**Supplemental Figure 8.**
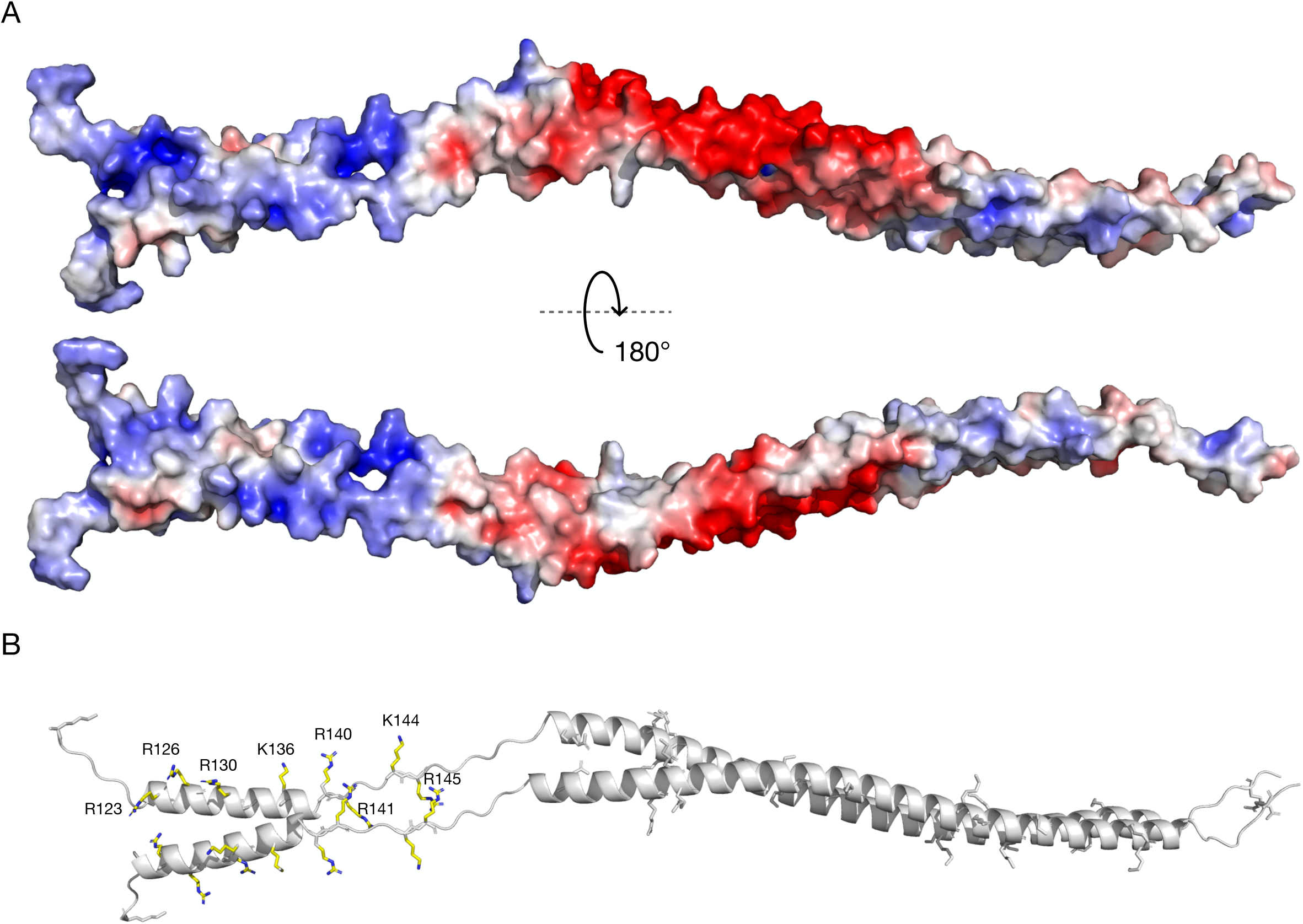
Lysine and Arginine distribution in the *T. brucei* KKT4^115–232^ model structure. (A) Homology model of KKT4^115–232^ coloured by surface electrostatic potential, showing positively charged patches in the N-terminus. Red to blue, −5 kbT to +5 kbT, as calculated by APBS electrostatic plugin in Pymol (Jurrus et al., 2018). (B) Lysine and arginine side chains are presented as sticks. Residues, which mutations affected the microtubule-binding activity of KKT4 are shown in yellow.

**Supplemental Figure 9.**
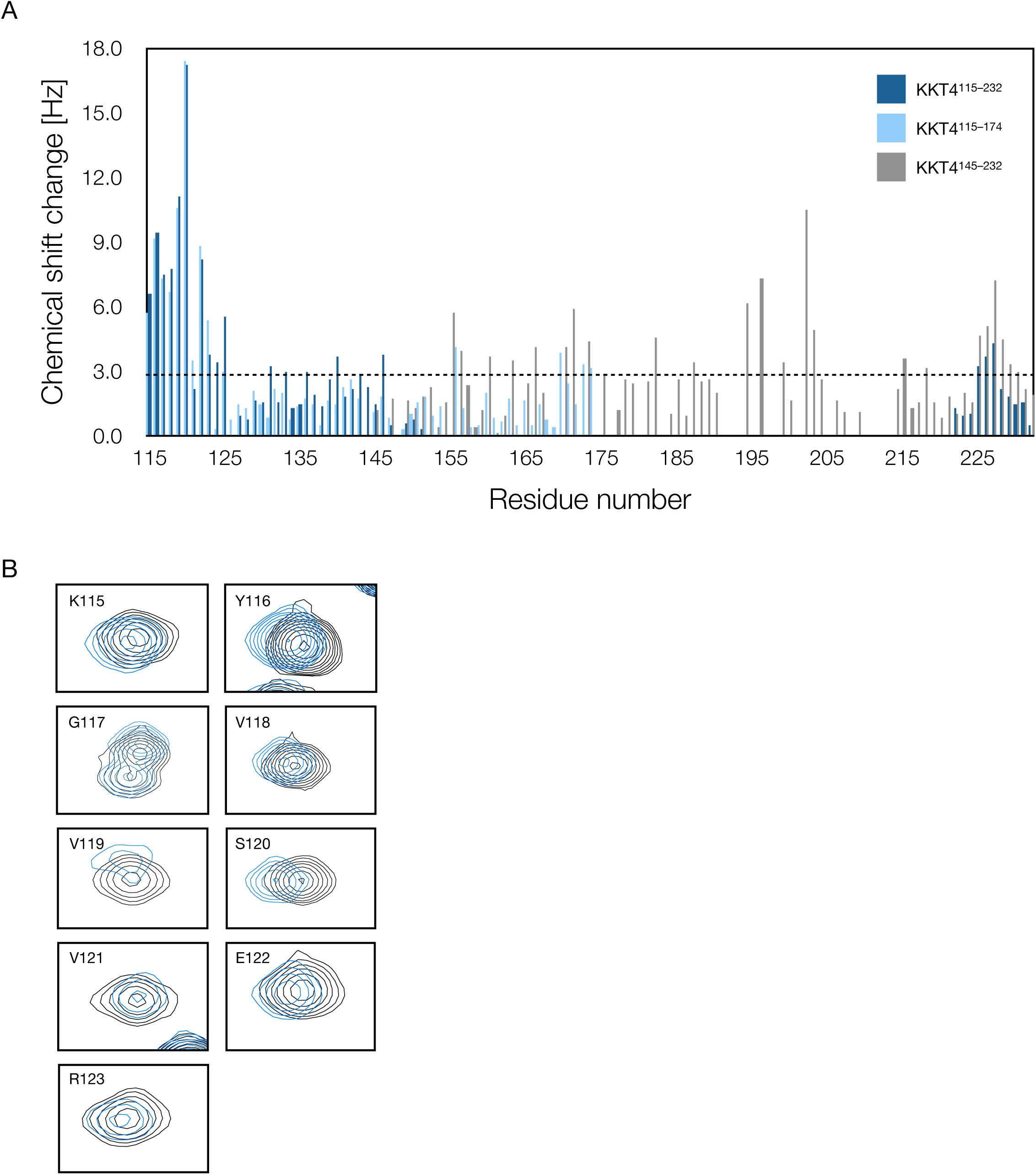
N-terminus of microtubule-binding domain directly interacts with BRCT domain. (A) Magnification of peaks that showed chemical shift changes upon addition of the KKT4 BRCT domain to the sample. Peaks coming from ^15^N-KKT4^115–232^ and ^15^N-KKT4^115–232^/KKT4^BRCT^ spectra are coloured in black and blue respectively. (B) Chemical shift changes observed upon addition of KKT4^BRCT^ to KKT4^115–232^ (dark blue), KKT4^115– 174^ (light blue), or KKT4^145–232^ (grey) have been plotted against the KKT4 sequence. The dotted line represents the average value (2.8 Hz) of all chemical shift changes observed.

